# Superinfection exclusion strategy of siphophage T5: analysis of the FhuA:Llp complex

**DOI:** 10.64898/2026.03.26.714494

**Authors:** Séraphine Degroux, Corinne Deniaud-Vivès, Emeline Metsdach, Claudine Darnault, Aline le Roy, Caroline Mas, Loïc Salmon, Torsten Herrmann, Cécile Breyton

**Affiliations:** Univ. Grenoble Alpes, CNRS, CEA, IBS, F-38000, Grenoble, France; Centre de Résonance Magnétique Nucléaire à Très Hauts Champs, UMR 5082, CNRS, Ecole Normale Supérieure de Lyon, Université Claude Bernard Lyon 1, Villeurbanne 69100, France; Univ. Grenoble Alpes, CNRS, CEA, EMBL, ISBG, F-38000, Grenoble, France

**Keywords:** Bacteriophage, Superinfection exclusion, induced fit interaction, two-step equilibrium, TonB dependent transporter

## Abstract

Superinfection exclusion is a widespread viral mechanism that protects the infected cell from over infection by other, identical or closely related viruses. Shortly after infection by bacteriophage T5, the bacterium *E. coli* produces Llp coded by the just injected viral genome. Llp is a small lipoprotein targeted to the inner-leaflet of the host outer-membrane. It binds to FhuA, an outer-membrane iron-ferrichrome transporter, which is T5 receptor at the *E. coli* cell surface. The interaction between Llp and FhuA prevents any further bacteriophage binding and infection. Here, we determined the RMN structure of Llp and analyse the formation of the FhuA:Llp complex using a wide range of techniques and mutants, both *in vivo* and *in vitro*. Interaction of Llp to FhuA is governed by a two-step equilibrium with a strong contribution of induced fit: Llp binding requires remodelling of FhuA periplasmic plug surface, allowing the further large conformational reorganisation of the extracellular loops. Analysis of FhuA mutants show the importance of the intertwined interactions between the extracellular loops and the plug in the communication between the periplasmic and the extracellular faces of FhuA.

## Introduction

When several viruses infect the same cell, there is superinfection. Superinfection allows genetical diversity in viral populations, is beneficial to environmental adaptation, but also enables defective virus to be maintained in the population (Hunter and Fusco, 2022). Superinfection exclusion (Sie) is the ability of a virus to block or interfere with a second infection by the same or a genetically related virus. Sie is a valuable short-term strategy, protecting viral multiplication (Hunter and Fusco, 2022). Sie has been described for all host virus systems, from human, animal and plant, to prokaryote (Broecker and Moelling, 2019). In bacteriophages – or phages – *i.e*. viruses infecting bacteria, several Sie systems have been described (Bucher and Czyż, 2024). For instance, in some *E. coli* prophages (phages whose sequences are inserted into the host chromosome), Cor proteins prevent superinfection by phages which receptor is the outer-membrane (OM) protein FhuA (Hernández-Sánchez et al., 2008). Although Cor proteins have been shown to functionally inactivate FhuA (Uc-Mass et al., 2004), the molecular mechanism responsible for this exclusion has not yet been elucidated. Regarding lytic phages, the Sie proteins Imm and Sp of the T4 lytic phage protect the host against T-even bacteriophage superinfection by blocking, either DNA ejection into the host cytoplasm at the inner membrane, or the peptidoglycan digestion, respectively (Lu et al 1994). The lytic conversion lipoprotein Llp of the lytic phage T5 (Braun et al., 1994; Decker et al., 1994) is another example of Sie protein, and is the object of the present work.

Phage T5 belongs to the *Caudoviricetes* order and to the siphophage paraphyletic morphological family; is known to infect *E. coli*. As other siphophages, T5 has an icosahedral capsid containing the genomic dsDNA and a long flexible tail. At its distal tip stand three L-shaped lateral fibres, and a straight fibre that bears the Receptor Binding Protein RBP_pb5_. The interaction of the lateral fibres with the OM lipopolysaccharides allows T5 to scan the bacterial surface (d’Acapito et al., 2025), ultimately leading to RBP_pb5_ binding to its receptor FhuA, an OM TonB-dependent ferrichrome transporter (Braun, 2009). Binding of RBP_pb5_ to FhuA commits T5 to infection, triggering the perforation of the bacterial cell wall and DNA ejection (Heller, 1984; Linares et al., 2023). The structure of the FhuA:RBP_pb5_ complex reveals RBP_pb5_ binding to FhuA (Degroux et al., 2023; van den Berg et al., 2022) and allowed to decipher the trigger of T5 infection (Degroux et al., 2023). After the initial phase of infection, phage T5 produces the Sie protein Llp that inactivates FhuA (Braun et al., 1994; Decker et al., 1994), protecting the bacterium turned into a phage factory. Llp main role is however probably to prevent the inactivation of progeny phages by their interaction with FhuA-containing cell debris (Decker et al., 1994). Recently, the structure of Llp:FhuA complex has been published (van den Berg et al., 2022).

The *llp* gene is located immediately upstream of the gene encoding RBP_pb5_, promoting their co-segregation by limiting genetic recombination. Llp, synthesised within 5 min post-infection, is a 7.09 kDa lipoprotein acylated on its N-terminal cysteine (Decker et al., 1994). Llp confers resistance towards all phages that use FhuA as a receptor. It is targeted to the inner leaflet of the *E. coli* OM (Decker et al., 1994; Robichon et al., 2003), where it can interact with the periplasmic surface of FhuA. FhuA:Llp interaction also prevents further transport of any ligand through FhuA, including its physiological ligand Fe-ferrichrome and the bacteriocin Colicin M (Braun et al., 1994; Decker et al., 1994). Llp was produced by recombinant expression and circular dichroism analysis revealed a β-structure fold (Pedruzzi et al., 1998). Llp possesses an internal disulphide bridge that is essential for its interaction with FhuA (Mondigler et al., 2006; Robichon et al., 2003). The FhuA:Llp structure confirms the periplasmic binding of Llp to FhuA and reveals rearrangement of two extracellular loops of FhuA explaining why RBP_pb5_ cannot bind to FhuA:Llp (van den Berg et al., 2022). Here, we determine the structure of free Llp by NMR, combine *in vivo* experiments, and biophysical and biochemical analyses, to comprehensively characterise the formation of the Llp-FhuA complex.

## Results

### 1. Biochemical characterisation of Ac-Llp and Sol-Llp

Production of full length acylated Llp (Ac-Llp), including its signal sequence, was tested in BL21(DE3) and C43(DE3). Both produced Ac-Llp, with a better yield in C43(DE3) (Fig. 1A). Ac-Llp expression in C43(DE3) at 37 °C with induction led to denser OM that contained non-solubilisable Ac-Llp aggregates. In contrast, expression at 20 °C without induction produced normal-behaving OM, containing Ac-Llp that could be solubilised and further purified. Under these conditions, the average purification yield was ∼3 mg of Ac-Llp per litre of culture. Mass spectrometry analysis of the purified Ac-Llp revealed four main species between 8737.9 and 8810.7 Da (Fig. S1A). Considering the theoretical mass of the his-tagged mature protein (7971.9 Da), these different masses are consistent with lipidation by a glycerol moiety bearing two fatty acids and an additional fatty acid linked to the terminal cysteine. The observed masses correspond to fatty acid chains ranging from C12 to C18, saturated or unsaturated, in agreement with the natural heterogeneity of fatty acid composition in *E. coli* membranes. The presence of a disulfide bond, required for interaction with FhuA (Mondigler et al., 2006; Robichon et al., 2003), cannot be confirmed here due to mass heterogeneity. Divalent cation binding to the protein was suggested (Pedruzzi et al., 1998), but when investigated by flame spectrometry, only traces of Ni^2+^ were found in our sample, probably due to a contamination from the Ni-NTA column purification.

**Figure 1.**
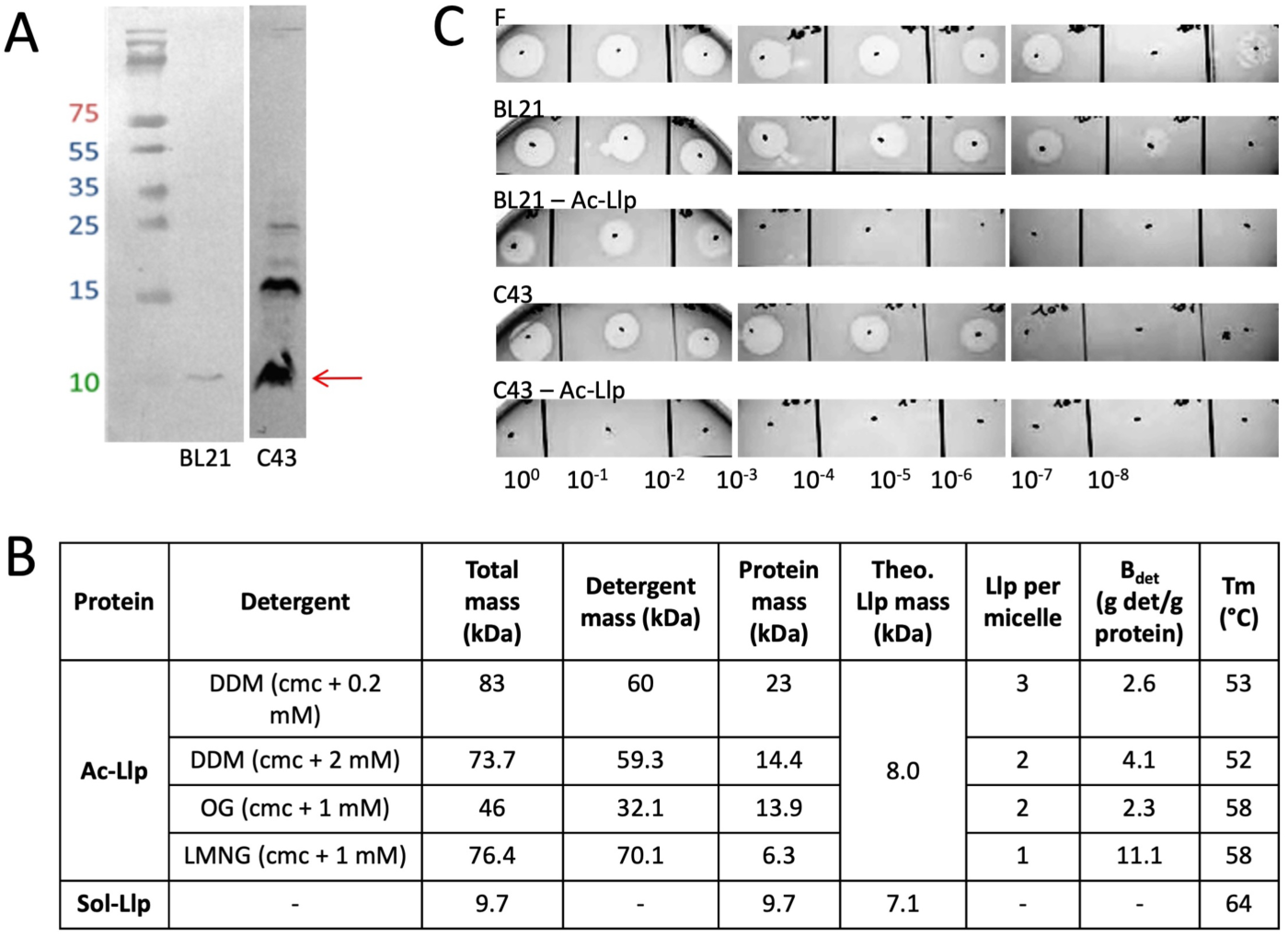
Functional and biochemical characterisation of Ac-Llp and Sol-Llp. **A.** Expression of Ac-Llp in whole-cell extracts, analysed by Western blot using an anti-His antibody. The Ac-Llp band is indicated with an arrow. The band at ∼15 kDa in the C43(DE3) sample is an artefactual dimer due to the presence of ß-mercapto-ethanol in the loading buffer. **B.** T5 sensitivity of strains F, C43(DE3), and BL21(DE3), either non-transformed or transformed with pET20b-Ac-Llp, was tested by spotting serial 10-fold dilutions of phage T5 (initial stock: 1.07 × 10¹³ PFU/mL) onto a bacterial mat. A decrease in detectable phage titre reflects an increased resistance to infection. **C.** Molar masses and oligomeric states and bound detergent (determined by SEC-MALS) and thermal stability (measured by nanoDSF) analyses of Ac-Llp in different detergent and of Sol-Llp.

The soluble form of Llp (Sol-Llp) was obtained as a fusion with MBP and removal of its N-terminal cysteine to prevent acylation. After MBP cleavage and purification, Sol-Llp was obtained with a yield of ∼20 mg per litre of TB culture. Mass spectrometry analysis revealed a mass of 7043.99 Da compatible with one disulphide bound. However, after incubation for a day at 30°C, MS data shows that ^15^N^13^C Sol-Llp loses four N-terminal amino acids. In contrast, Ac-Llp N-terminal is not degraded over time, as it is probably protected/stabilised by the acylation and the bound detergent micelle. The maximum thermal stability for Ac-Llp in DDM and Sol-Llp is observed between pH 5.6 and 6.6 with a Tm of 53°C and 64°C, respectively, and Ac-Llp is more stable when transferred in LMNG and OG (Tm = 58°C) (Fig. 1B). The presence of a disulphide bridge in both proteins was confirmed by a 6.5°C decrease of their respective Tm in the presence of a reducing agent.

The oligomerisation state of the two proteins was investigated by SEC-MALLS: Sol-Llp is predominantly monomeric, with only minor amounts of dimer. For Ac-Llp, SEC-MALLS also allowed to measure the amount of bound detergent (B_det_, in g of detergent/g of protein). Results for Ac-Llp in 0.4 mM DDM (cmc (critical micellar concentration) + 0.2 mM) suggest that there are three Ac-Llp molecules per DDM micelle with a B_det_ of 2.6 g/g, whereas in 2.2 mM DDM (cmc + 2 mM), the analysis gives two Ac-Llp per micelle with a B_det_ of 4.1 g/g. A similar result was observed in 21 mM OG (cmc + 1 mM), where the small free OG micelles are separated from the Ac-Llp-detergent complex. In contrast, in 1 mM LMNG (cmc + 1 mM), results are consistent with only one Ac-Llp per micelle with a B_det_ of 11.1 g/g (Fig. 1B). This large amount is compatible with the propensity of LMNG to form long cylindrical micelles (Breyton et al., 2019).

### 2. Sol-Llp and Ac-Llp share the same structure

Ac-Llp in DDM and Sol-Llp CD spectra were measured to validate the global fold of Sol-Llp prior to NMR structure determination. As previously published (Pedruzzi et al., 1998), the CD spectra of the two proteins are very similar (Fig. 2A). There is a small difference at around 230 nm probably due to the N-terminal of Ac-Llp that could interact with the detergent micelle. The predictions of the secondary structure proportions confirm the visual aspect of the curves and suggest a β-sheet content of 40% in both proteins. Contrary to what is presented in Pedruzzi *et al*., EDTA does not affect the CD spectra of either proteins. To complement these results, we compared the TROSY-methyl-TROSY spectra of Ac-Llp and Sol-Llp recorded in natural abundance of ^13^C. To allow better relaxation times, Ac-Llp was transferred into OG, which forms smaller micelles than DDM. As expected, the spectrum of detergent-solubilised Ac-Llp remains much less intense than that of Sol-Llp and is dominated by the signal of the detergent aliphatic chains. Nonetheless, the rest of the spectra is very similar to that of Sol-Llp (Fig. 2B), confirming that the global structures of Sol-Llp and Ac-Llp are similar.

**Figure 2.**
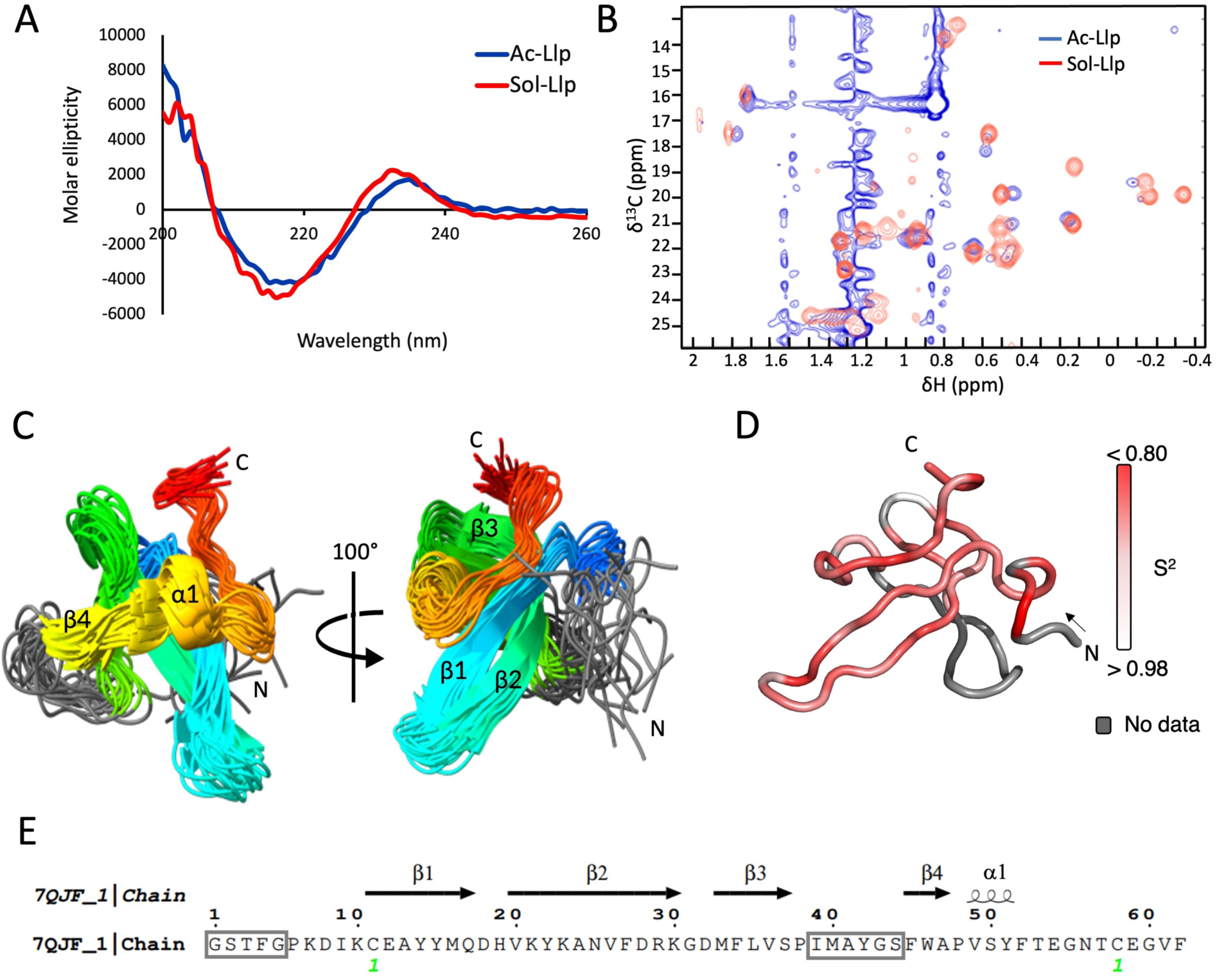
Structural fingerprints and backbone dynamics of Sol- and Ac-Llp. **A.** Circular dichroism spectra of Ac-Llp solubilised in DDM (blue) and Sol-Llp (red). Each curve represents the average of 10 acquisitions, corrected by subtraction of buffer spectra (10 averaged scans) and normalised to protein concentration. **B.** TROSY-methyl-HSQC spectra of Ac-Llp solubilised in OG (blue) and Sol-Llp (red). Spectra were recorded overnight at natural ^13^C abundance on a 600 MHz spectrometer, using protein concentrations of 0.91 mM (Sol-Llp) and 1.15 mM (Ac-Llp) in 25 mM Tris buffer, pH 6.5 at 303°K. The signal of the three methionines of the protein are easily identifiable in the 2 – 1.8 ppm area. The aliphatic region of the Ac-Llp spectrum (1.6 – 0.8 ppm) shows additional peaks arising from detergent aliphatic chains. **C.** NMR structure of Sol-Llp. Ensemble of the 20 lowest-energy conformers, coloured from the N-terminus (blue) to the C-terminus (red). Atomic coordinates are well defined except for residues 1–5 and the 39–44 loop (coloured in grey). Secondary structure elements (β1–β4 and α1) are indicated according to the assignment derived from ESPript analysis. **D.** Model-free analysis of Sol-Llp backbone dynamics. Order parameters (S²) are mapped onto the protein structure, from white (rigid) to bright red (mobile). Residues without data are shown in grey. **E.** Linear representation of Sol-Llp secondary structure as extracted from the NMR structure (PDB ID 7QJF). β-strands (β1–β4) and the short 3₁₀-helix (α1) are indicated above the sequence. The cysteine residues involved in the disulfide bond are labelled green. Non-assigned amino acids in NMR are boxed in grey.

### 3. Structure of Sol-Llp

The NMR study was focused on Sol-Llp. A set of NMR experiments were carried out to allow spin system assignment and structure calculations. Spin system assignment from 3D NMR spectra (using CCPNMR3) covered 47 residues – 76% of the Sol-Llp sequence. Weaker additional correlations indicated a minor conformational subpopulation likely linked to N-terminal degradation. A total of 511 meaningful, non-redundant upper Nuclear Overhauser Enhancement (NOE)-derived distance restraints were automatically generated by the NMR analysis program UNIO (Guerry et al. 2015) in combination with the NMR structure calculation program CYANA (Güntert et al., 1997). The final NMR structure bundle was calculated in explicit solvent using the program CNS (Brunger, 2007). The 20 NMR conformers have very good stereochemistry with 95.2% of the residues in most favoured regions and additionally allowed regions. In addition, the structural ensemble exhibits excellent geometry with no systematic violation of distance restraints greater than 0.2 Å and no systematic violation of dihedral angles greater than 5°. With the exception of the first five N-terminal residues and the 39-44 loop, the atomic coordinates of the NMR structure are well-defined. The root-mean-square-deviation (RMSD) value from the mean coordinate positions of the backbone atoms for residues 6–38 and 45–62 is 1.02 ± 0.19 Å (Fig. 2C). For the regions 1-5 and 39-44, peaks can be attributed to the backbone, but no NOE can be identified. Detailed NMR structure and energy refinement statistics are reported in Table S1.

Sol-Llp adopts a compact β-rich fold composed of a short helical segment (α1) and four β-strands (β1–β4) organised into two β-sheets, connected by an unresolved loop. The N-terminal is linked to the C-terminal by a disulfide bond between residues C11 and C58 (Fig. 2C,E). Backbone dynamics of Sol-Llp was investigated using NMR ^15^N spin relaxation to probe internal motions at the ps to ns timescales. Longitudinal (R_1_) and transverse (R_2_) relaxation and heteronuclear NOEs were measured to investigate backbone flexibility. Using a model-free analysis, it was possible to determine a global rotational correlation time of 6.3 ns, in agreement with a monomeric protein of this size and investigate local motions. Overall Llp is quite rigid with a high average order parameter (S² ≈ 0.88), consistent with limited backbone motions across most of the protein (Fig. 2D, S2). Despite this global rigidity, local dynamics were detected: H19, Y22, A24 and M33 showed elevated R_1ρ_ values, indicative of conformational exchange. Increased flexibility appears towards the N- and in a lesser proportion, towards the C-termini. Moreover, the large loop spanning residues 39–44 could not be assigned in the 3D spectra and displayed no long-range NOEs, both features being typical of flexible/disordered regions. Taken together, these data indicate that Sol-Llp is globally rigid but harbours localised dynamic regions, including a mobile loop (residues 39–44) that may play a functional role.

### 4. T5 protection of cells overexpressing Llp

T5 sensitivity tests were used to assess Ac-Llp functionality *in vivo*. BL21(DE3) showed similar T5 sensitivity to the reference F strain, whereas C43(DE3) was less sensitive, with a two-log reduction in T5 titre (Fig. 1C). Upon transformation with the *Ac-Llp* plasmid and overexpression (induced with IPTG) of the Ac-Llp protein, C43(DE3) exhibited higher protein levels than BL21(DE3), as checked by Western blot (Fig. 1A). This is well correlated with the complete protection of C43(DE3) expressing Ac-Llp against T5, while BL21(DE3) expressing Ac-Llp was only partially protected. To examine the impact of the FhuA:Llp ratio on *in vivo* protection, we modulated protein expression levels independently, taking advantage of the inducible or leaky expression of Ac-Llp and the endogenous, iron-regulated expression of FhuA in BL21(DE3) (Fig. S3). When both FhuA and Ac-Llp were uninduced (in the absence of Dipyridyl and IPTG, respectively), cells were partially protected with a 5-log reduction compared to the Llp empty strain. Overexpression of Ac-Llp alone increased protection (7-log reduction) and induction of FhuA alone provided only low protection (2-log reduction). Induction of both proteins did not increase protection (5-log protection, as for the uninduced situation). The C43(DE3) strain was protected whatever the condition, due to a strong expression of Llp even without induction (Fig. S3C). Thus T5 resistance *in vivo* depends on the ratio between FhuA and Llp rather than on their total amounts.

### 5. Ac-Llp forms a stable and functional complex with FhuA

We then checked whether the Ac-Llp:FhuA complex is functional *in vitro* by investigating whether the FhuA:Llp complex could “protect” T5. T5 was incubated with either FhuA or Ac-Llp:FhuA, and T5 infectivity then was monitored on a sensitive *E. coli* strain (Fig. S4A). When incubated with FhuA, T5 is inactivated for subsequent infection. If FhuA is pre-incubated with Llp forming a functional complex, it can no longer interact with T5, and T5 remains infectious (Fig. S4B). Different FhuA:Ac-Llp stoichiometries were tested, with pre-incubation at 4, 20 or 37°C. At 4°C, a functional complex was obtained at 1:1 and 1:8 FhuA:Ac-Llp ratio after 96 h and 48 h pre-incubation, respectively (Fig. S4C). At 20°C and with 1 h pre-incubation, only the 1:130 and 1:200 ratios efficiently protected T5. When pre-incubated at 37°C, the 1:8 ratio provided partial protection after 30 min and complete protection after 4 h; the 1:1 ratio started to protect T5 after 2 h of pre-incubation. Sol-Llp could not form a stable, functional complex with FhuA, neither at FhuA:Sol-Llp 1:30,000 without incubation, at room temperature, nor at 1:40, pre-incubated seven days at 4°C. Thus, FhuA:Ac-Llp forms a functional complex, but it requires a high activation energy (either long pre-incubation time or high temperature).

### 6. Sol-Llp and Ac-Llp interact *in vitro* with FhuA

Several techniques were considered to measure the K_d_ of the Sol-Llp:FhuA interaction. SPR and BLI analyses were carried out, but the detergent necessary to solubilise FhuA induced a large signal bias that prevented any statistical analysis and no reliable K_d_ could be determined. No interaction could be detected by SEC-MALLS nor AUC, even after pre-incubation (Fig. S5A). Only MST gave clear results, with a K_d_ of 1.5 ± 0.3 mM (Fig. 3A). An NMR titration, adding ^15^N-Sol-Llp to unlabelled-FhuA was performed, with the acquisition of a [^1^H-^15^N] SOFAST-HMQC spectrum for each titration point. The maximal fraction of Llp engaged in the complex is limited by the initial concentrations of FhuA (0.15 mM) and Llp (1.2 mM). Considering the measured K_d_, only ∼8% of Llp is expected to be FhuA-bound at the highest Llp concentration (Fig. S5B). Nonetheless, the clear variations in the chemical shifts and peak intensities are consistent with a K_d_ in the mM range. Further analysis of these experiments for defining the interaction surface of Llp with FhuA are given below.

**Figure 3.**
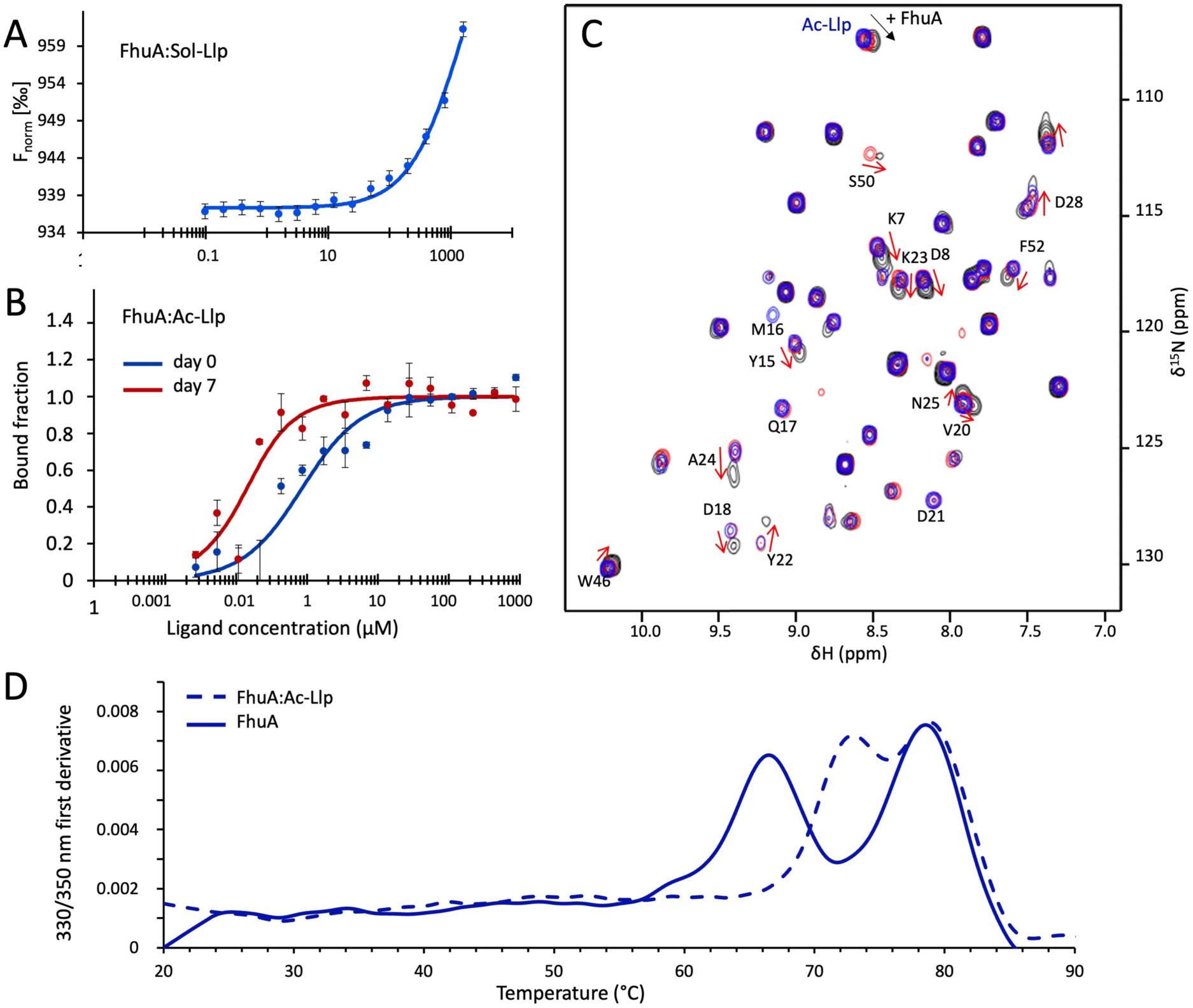
Characterisation of the FhuA:Llp complex. MST affinity curves of Sol- (**A**) and Ac-Llp (**B**) titrated against fluorescently labelled FhuA (125 nM). Sol-Llp was tested in a concentration range from 24 nM to 1.6 mM and Ac-Llp from 0.027 to 887.2 µM, both in 25 mM Tris, pH 8.5, 150 mM NaCl and 0.2 mM DDM. For the Ac-Llp-FhuA sample, MST was performed either immediately after mixing the proteins (red) of after pre-incubation of seven days at 4°C (blue). **C.** [^1^H,^15^N] SOFAST-HSQC spectra of ^15^N-labeled Sol-Llp in DDM during titration with detergent-solubilised FhuA. The spectrum of Sol-Llp in DDM is shown in blue. Addition of FhuA induces progressive chemical shift perturbations and signal intensity changes, consistent with complex formation. Red: 46 µM Sol-Llp, 127 µM FhuA, (0.4% complex for a K_d_ of 1.5 mM). Black: 436 µM Sol-Llp, 59 µM FhuA (7.4% complex). Red arrows indicate residues with significant chemical shift perturbation label. Labelled residues without an arrow indicated loss of intensity signal throughout the titration. **D.** Thermal denaturation curves of FhuA (solid line) and of FhuA:Ac-Llp after two days incubation at room temperature (dashed line) measured by NanoDSF.

Concerning the Ac-Llp:FhuA complex, we checked first whether the formation of the complex affected FhuA stability. Thermal unfolding profile of FhuA presents two denaturation transitions corresponding to that of its plug and of its 22-strand β-barrel (Fig. 3D). With prior incubation two days at room temperature with Ac-Llp, FhuA plug denaturation transition is shifted to higher temperatures, the Tm increasing from 65.8°C to 72.8°C, indicating that the Ac-Llp:FhuA interaction stabilises FhuA plug (Fig. 3D). Different methods were considered to determine Ac-Llp:FhuA K_d_. SPR and BLI experiments were here again imprecise due to detergent artefacts, but interactions were however detected between Ac-Llp and FhuA regardless of the protein bound to the surface, and resulted in K_d_ estimates ranging from 0.5 to 50 µM. To avoid detergent artefacts, SV-AUC was considered. Results were however not reproducible and occasionally SV-AUC did not even show formation of the Ac-Llp:FhuA complex. Nevertheless, when FhuA and Ac-Llp were pre-incubated for 7 days at 4°C before analysis, a complex was systematically observed, with a shift in the sedimentation coefficient from 7.8 to 8 S (Fig. S6). Different K_d_ values were also determined by MST, whether measured immediately after sample preparation at room temperature, leading to a K_d_ of 1 µM ± 0.3 or after seven days pre-incubation at 4°C, leading to a K_d_ of 0.1 ± 0.05 µM (Fig. 3B).

Thus, Sol-Llp and Ac-Llp interact with FhuA in the mM and µM range, respectively, indicating that acylation is essential to form a stable and functional complex between Llp and FhuA. The Ac-Llp:FhuA interaction involves two equilibria with K_d_ differing by one order of magnitude.

### 7. Llp interaction surface with FhuA probed by mutations in Llp

The SOFAST HMQC RMN spectra of ^15^N-Sol-Llp incubated with unlabelled FhuA showed chemical shifts and/or intensity perturbations for 33 residues over 47 attributed residues, *i.e*. 70% of the protein (Fig. 3C). We designed a variety of mutations, from point mutations to broader “patch” mutations covering neighbouring residues intended to disrupt the interaction: D28K, KD7-8AR, YK22-23AS, RK29-30SS, YYHQD14-18AFAK, D18K21K23-NNQ and FTE52-54YKK. In order to probe the flexible loop, which is not visible in NMR, W46S, as well as a larger cluster, YGSFW42-46STRAS, were designed (Table 1, Fig. 4). The C43(DE3) strain was transformed with these mutated *Ac-Llp* plasmids and tested for protection from T5. In the following, log differences refer to reductions in protection relative to the maximal protection conferred by WT Ac-Llp. Mutants could be classified into 4 groups: mutant KD7-8AR behaved as the WT protein, protecting the bacteria against T5 infection; mutants YK22-23AS, RK29-30SS and W46S retained strong partial protection, with only a minor decrease protection (∼2-log). D28K, FTE52-54YKK and D18K21K23-NNQ mutants showed weak partial protection against T5 infection (3-4 log), suggesting Llp is not able to bind efficiently FhuA (Fig. 4A). All mutants reached expression levels that should be sufficient for full protection in C43(DE3) (Fig. 4B).

**Table 1:**
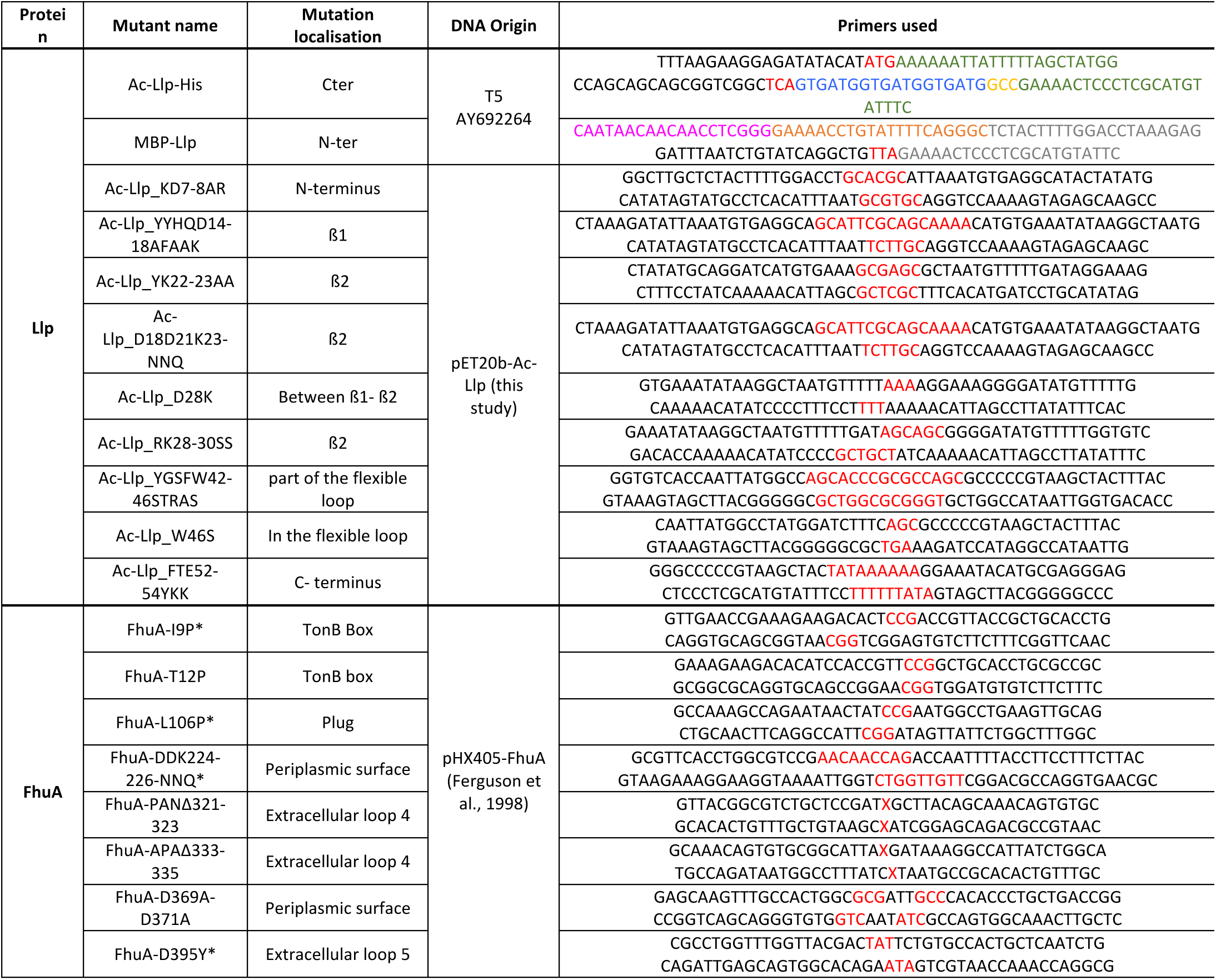
Summary of Llp and FhuA mutants with the corresponding primers and mutation location. FhuA is numbered without the His-tag inserted in loop 4, and Llp is numbered without the signal sequence. * Mutants repeated from Braun et al., 1994.

**Figure 4:**
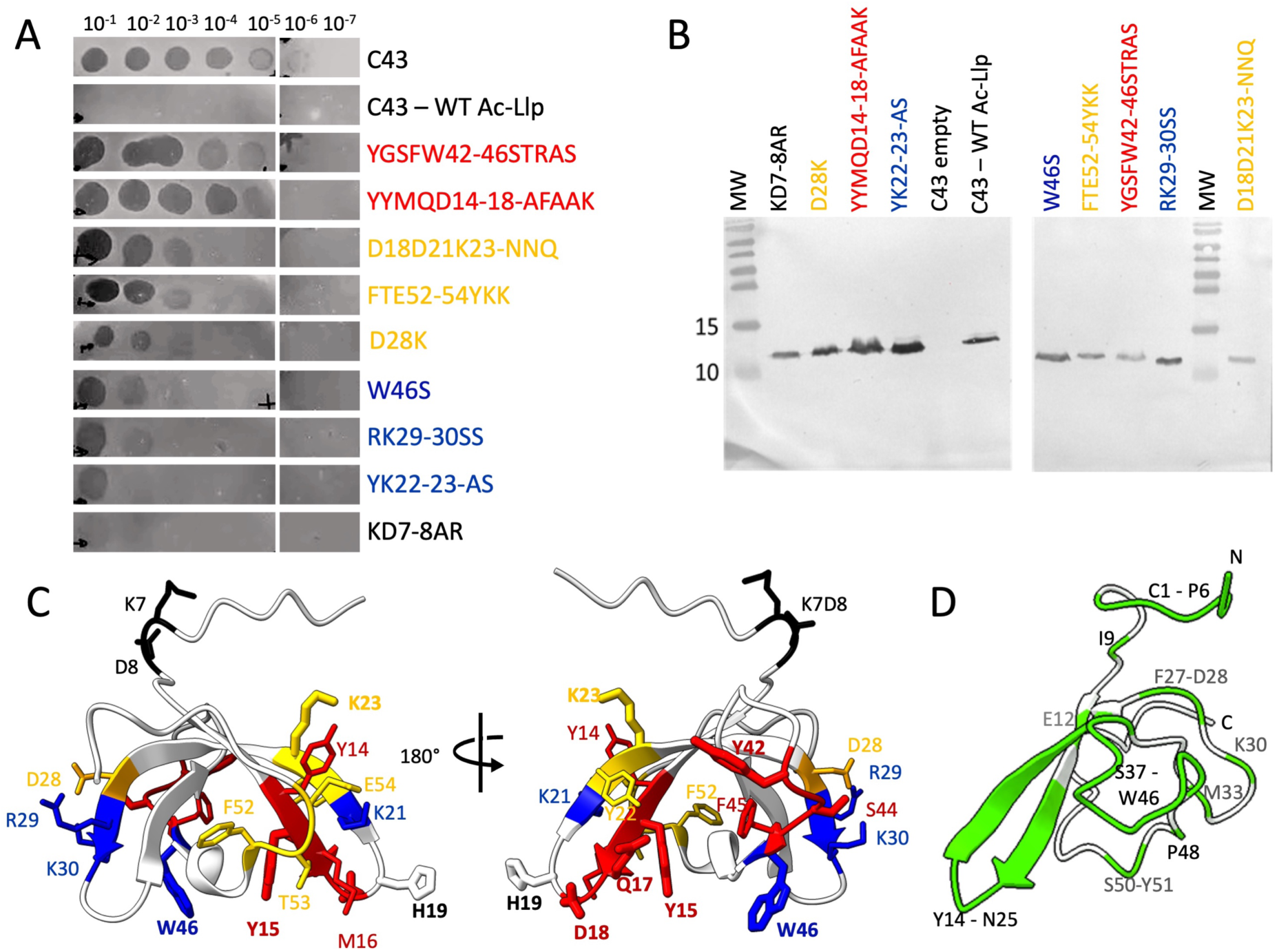
Analysis of Ac-Llp mutants. **A.** Bacterial lawns of *E. coli* C43(DE3) expressing WT or mutant Ac-Llp were spotted with serial dilutions of phage T5. WT Ac-Llp confers full T5 resistance to the C43(DE3) strain, whereas mutants display a continuum of phenotypes grouped into four classes: black as WT; blue, partial protection; yellow, weak protection; red, no protection. The same colour code is used in panels B and C. **B.** Western blot analysis of total C43(SE3) cell extracts using an anti-His antibody shows that all Ac-Llp variants are expressed without induction, with moderate variations. These differences do not correlate with the protection phenotype observed in panel A. **C.** Mutated residues are shown in sticks and colour-coded according to their T5 protection phenotype (same colour code as in panel A) on the Sol-Llp ribbon structure. Side chains of residues forming hydrogen bonds or salt bridges with FhuA in (van den Berg et al., 2022) are shown in thicker sticks to highlight their functional importance. **D.** Residues involved in the interaction interface with FhuA are coloured in green and labelled on the ribbon representation of FhuA-bound Ac-Llp (PDB 8A60). Black labels residues in the front, grey in the back.

### 8. FhUA interaction surface with Llp probed by mutations in FhuA

Braun *et al.,* (1994) described FhuA mutants, distributed over the protein, which were insensitive to Llp-mediated protection. We reproduced some of these mutants and others to determine whether the loss of protection resulted from impaired Llp binding or deficient signal transduction within FhuA (Fig. 5A). Mutants include Braun’s point mutants I9P (tonB box), D395Y (loop EL5) and L106P (plug); we produced smaller deletions than that of Braun: APAΔ333-335 and PANΔ321-323 (loop EL4). We introduced D369D371-AA and DDK224–226NNQ substitutions in the periplasmic turns T2 and T10 to test the role of these charged residues, and the T12P substitution (TonB box) to evaluate the requirement of TonB interaction for Llp activity. This latter mutant is impaired in transport, as it is resistant to Colicin M.

**Figure 5:**
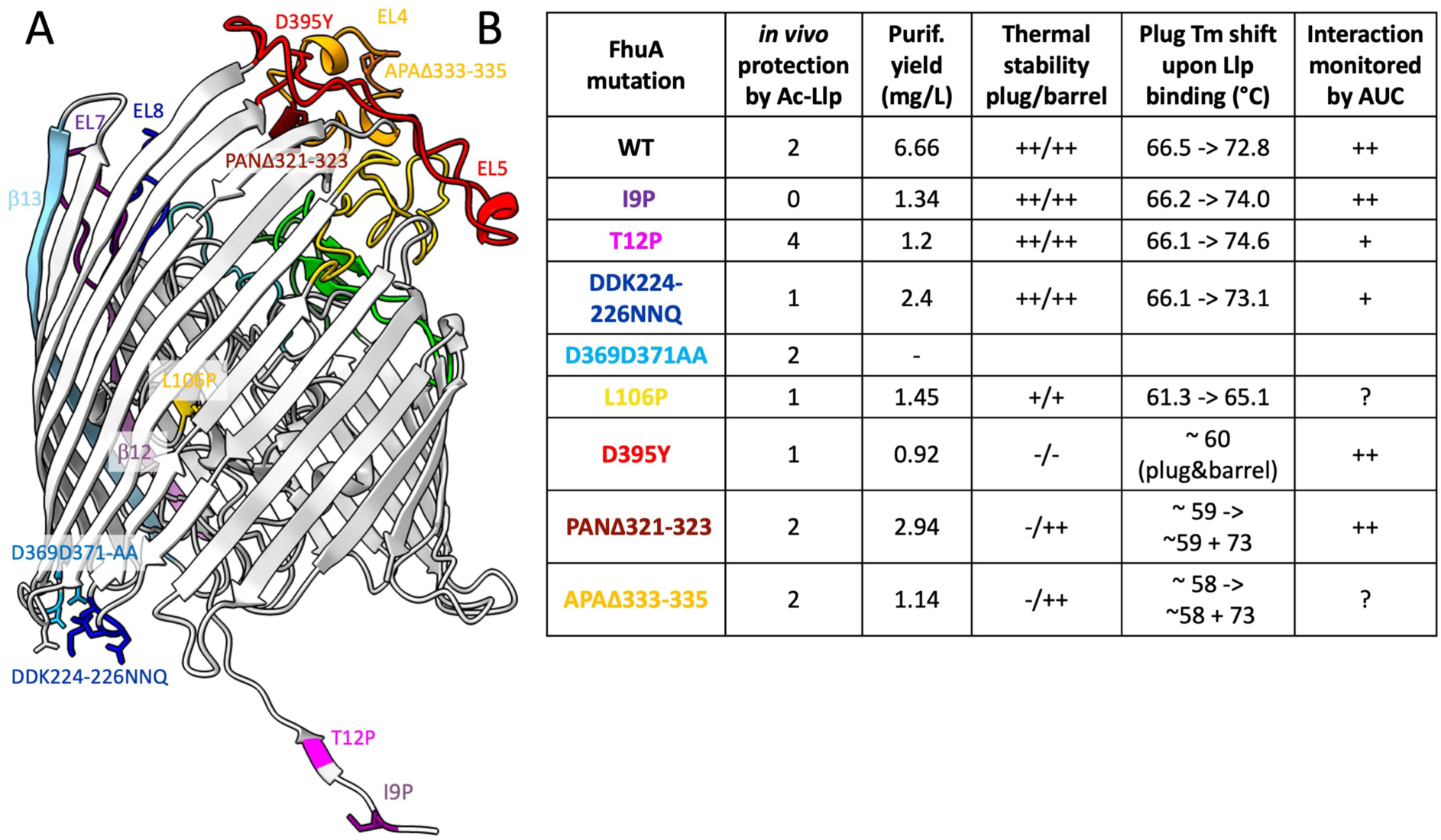
Analysis of FhuA mutants. **A.** Mutated residues are represented as sticks on the ribbon structure of FhuA in complex with TonB C-terminus (PDB 2GRX). Extracellular loops are coloured, and EL4, 5, 7 and 8 are labelled, as well as β-strands 12 (plum) and 13 (light blue), showing barrel discontinuity its upper part. **B.** Summary of the biochemical and biophysical properties of the FhuA mutants. *In vivo* Ac-Llp protection by Ac-Llp on AW740 expressing the different FhuA mutants and Ac-Llp: the number represents the log reduction in phage titre relative to the control without Ac-Llp. Thermal stability of FhuA plug and barrel: ++, WT Tm values; +, lower but clear transition; -, unclear transition. Interaction monitored by AUC: ++, clear FhuA sedimentation shift upon incubation with increasing Ac-Llp concentrations; +, less clear shift; ?, uncertain shift.

The *E. coli* AW740 strain, deleted for the endogenous *fhua* gene and historically used for FhuA overexpression (Ferguson et al., 1998), was first transformed with the plasmids encoding the different FhuA mutants under the control of FhuA native promoter. Except for mutant D369D371-AA, all mutants were expressed approximately to the same level in the absence of induction by iron depletion, and all strains were sensitive to T5 at the same titre. The different strains were then co-transformed with an *Ac-Llp* encoding plasmid under an anhydrotetracyclin-inducible promoter, and again challenged with T5. Ac-Llp expression is expected to result in a protection of *E. coli* toward T5 infection. Even though anhydrotetracyclin was present and dipyridyl absent at all steps, *i.e.* with induction of Ac-Llp only, the strain co-expressing WT FhuA and Ac-Llp was only poorly protected. This could be due to an expression level difference of FhuA and Ac-Llp or to the absence of Omps in the AW740 strain (Arguijo-Hernández et al., 2018). Nevertheless, when comparing the T5 titres, FhuA mutants were classified into three categories. In the following, log differences refer to the absolute level of protection against phage T5. The reduction in T5 phage titre relative to the control without Llp is 2 logs for the strain expressing WT FhuA. Cells expressing FhuA mutant I9P (TonB box) showed no reduction in titre: it is unprotected by Ac-Llp co-expression. Cells expressing FhuA mutants L106P (plug), D395Y (EL5) and DDK224–226NNQ (T10) exhibited a 1-log reduction in apparent phage titre: they are partially protected by Ac-Llp co-expression. Finally, the cells expressing the remaining FhuA mutants (in T2, EL4) showed a 2-log reduction titre, *i.e.* they are protected by Ac-Llp similarly to the WT, and mutant T12P showed the highest level of protection, corresponding to a 4-log decrease in phage titre (Fig. 5B). Mutants reproduced from Braun et al., gave the same results.

The purification yield of the mutated FhuA ranged between 2 and 7 times less than that of the WT protein despite induction by iron depletion (Fig 5B). FhuA:Ac-Llp complexes of all purified FhuA mutant proteins were analysed by nano-DSF, after two weeks incubation at 4°C. The Tm of the plug is higher in the presence of Ac-Llp for FhuA mutants I9P, T12P, DDK224-226-NNQ and L106P, as for WT FhuA, suggesting that the FhuA:Ac-Llp complex forms (Fig. S7A). For FhuA mutants D395Y, PANΔ321–323, and APAΔ333–335, the stability of the plug appears compromised by the mutations in unbound FhuA, precluding definitive conclusions regarding their interaction with Llp. However, a small shoulder at the position of the Llp-bound plug in the WT FhuA is present in the two deletion mutants, suggesting that when Llp is bound, the plug assumes the WT conformation (arrow in Fig. S7). This analysis was completed by SV-AUC study of the mutated FhuA:Ac-Llp complexes, after seven days incubation at 4°C. SV-AUC confirmed an interaction between FhuA and Ac-Llp for FhuA mutants I9P, D395Y and PANΔ321-323, with an increase of ∼0.2 S of the sedimentation coefficient of FhuA. For FhuA mutants T12P and DDK224-226-NNQ, AUC showed a weaker interaction. For FhuA mutants L106P and APAΔ333-335, the results remained unclear. MST was performed only for FhuA mutants I9P and T12P, and show a similar interaction than for the WT. Results are summarised in Fig. 5B. Taken together, these data show that the absence of protection of the mutants investigated by Braun et al. does not result from defective Llp binding but rather from a defective signal transduction within FhuA (see further discussion below).

## Discussion

### 1. Cor proteins and Llp

Phage proteins are known to exhibit high sequence divergence, often rendering sequence-based homology detection tools ineffective. However, despite this variability, they frequently retain conserved structural folds (Veesler and Cambillau, 2011; Linares et al., 2020). To investigate whether this principle holds for the Cor and Llp families of Sie proteins, which both target FhuA, we predicted the structures of the mature form for various Cor and Llp proteins identified in temperate and lytic phages. Structural predictions of the Cor/Llp proteins encoded by phages T5, T1, φ80, mEp167 and HK022 revealed high-confidence models with similar structural architecture (RMSD values ranging from 6.4 to 7.5 Å over the full-length proteins), despite low sequence identity (18 to 25%). The main differences concern the 39-44 loop and the N-terminus, which are flexible elements of Llp (Fig. S8). Overall, it confirms that Llp and the Cor proteins belong to the same family of proteins.

When the structure of Llp is submitted to DALI to search for proteins of known or predicted structure with similar fold (Holm, 2020), a phage associated cell-wall hydrolase and three phage endolysins are identified. Llp aligns only partially on their larger CHAP (cysteine, histidine-dependent amidohydrolases/peptidases) domain, with poor Z-scores ranging from 3.0 to 3.3. A DALI search on AFViroColabFold does not retrieve any hit, and AF-Ecoli retrieves three irrelevant hits, which is surprising given the structural similarity of Llp with the Cor proteins of prophages. A FoldSeek search (van Kempen et al., 2024) on the other hand hits many small phage proteins in the Big Fantastic Virus Database (BFVD)(Kim et al., 2025), suggesting that the Llp fold is a common one among phages and probably a common Sie strategy to target TonB-dependant outer-membrane transporters.

### 2. Allosteric inhibition of FhuA by Llp and analysis of FhuA mutants

The crystallographic structure of the FhuA:Llp complex revealed that Llp binding triggers conformational rearrangements in the plug of FhuA. More specifically, the movement of loops EL7 and EL8 blocks access to the receptor and thereby prevents phage infection (van den Berg et al., 2022). We analyse our FhuA mutants in the light of this structure: a first group of mutations include D395Y (loop EL5), PANΔ321–323 and APAΔ333–335 (loop EL4). NanoDSF thermal unfolding profiles of these FhuA mutants show a poorly defined plug transition, with only a small fraction of plug stabilisation upon Llp incubation (arrow in Fig. S7A), suggesting a poor stability/improper folding of the plug. Interestingly, these mutations are located in the extracellular loops, far from the plug. Structural inspection of FhuA shows an intertwined connection between the different loops, ultimately interacting with the tip of the plug: in particular D395 contributes to EL5 stabilisation through interaction with S326 and Q328 (EL4). The behaviour of our FhuA mutants shows that these intertwined interactions are critical to stabilise the plug and that small changes as those introduced by mutations in EL4 and EL5 compromise the structural integrity of the plug. SV-AUC analysis of these FhuA mutants:Ac-Llp complexes indicate however that Ac-Llp is still able to bind FhuA mutants D395Y and PANΔ321–323. It suggests that plug correct folding or stability is not required for Ac-Llp binding. This is coherent with the FhuA:Ac-Llp structure, which shows that Ac-Llp interacts with more residues from the barrel than from the plug. On a functional point of view however, FhuA mutant D395Y (EL5) is less protected by Ac-Llp, whereas the mutants APAΔ333-335 and PANΔ321-323 (EL4) behave as the WT protein. We conclude that the plug is more severely destabilised in FhuA mutant D395Y than in the two deletion mutants, in which it can still undergo the conformational rearrangements necessary to trigger loops EL7 and EL8 repositioning when Ac-Llp is bound, as suggested by the shoulder in the nanoDSF curve (arrow in Fig. S7). The analysis of FhuA mutants confirms that the Llp binding information is transmitted to the extracellular loops through conformational changes of the plug.

The rearrangements of the plug are triggered by the remodelling of its periplasmic surface: residues 20-51 of the Apo FhuA (dark green in Fig. 6A,B), are displaced in the Llp-bound form and point out, partially disordered, into the periplasm (deep pink in Fig. 6A,B). This displacement allows Llp to “penetrate” into the barrel, thereby further stabilising the plug (as indicated by the increase of its Tm upon Llp binding). This remodelling allows a downward shift of loops 134-146 then 95-103 (red arrows, Fig. 6B). The downwards shift of FhuA plug loop 95-103 in turn allows subtle movements within the β-barrel wall, ultimately leading to the collapse of EL7 and EL8. Indeed, because β-strands 12 and 13 of the barrel are structurally independent from the rest of the barrel in its upper part (black double arrow in Fig. 6B), strand 13, and with it EL7 and EL8 can shift downwards (orange arrows, Fig. 6B). The L106P mutation, located near loop 95-103 and conferring to the protein only partial protection by Llp binding, likely impairs correct folding of the plug, and thus the concerted plug-barrel movement which is required for Llp-induced T5 protection.

**Figure 6:**
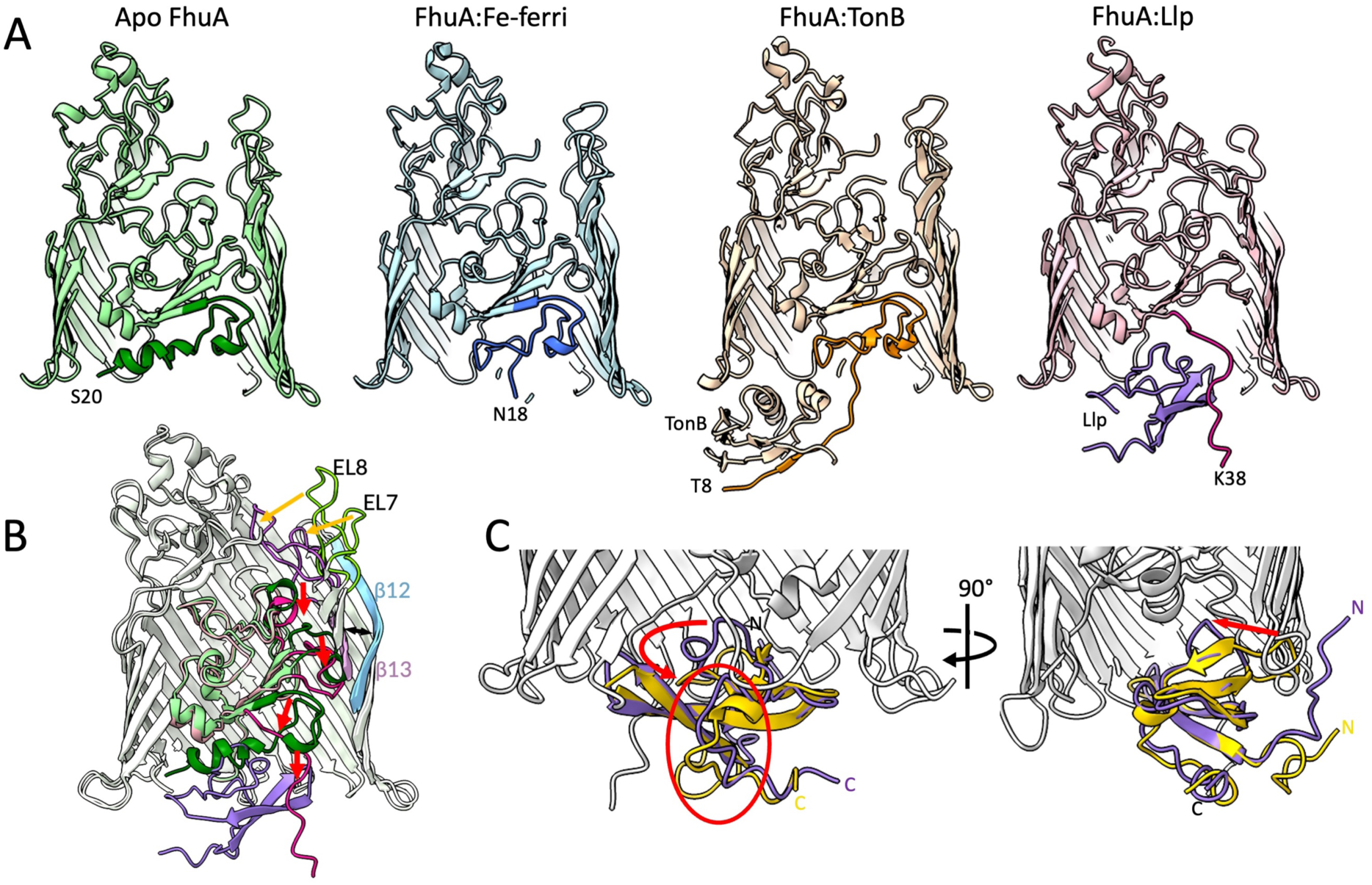
Comparison of FhuA and Llp structures in different states. **A.** From left to right: apo FhuA (PDB 1BY3, green), FhuA:Fe-ferrichrome (not represented)(PDB 1BY5, blue), FhuA:TonB (PDB 2GRX, orange) and FhuA:Llp (PDB 8A60, pink, Llp purple). The comparison shows the restructuration of the N-terminus that forms the plug periplasmic surface (coloured in darker shade until residue 51). The first N-terminal resolved residue is labelled. Front and back strands of the FhuA barrel have been sliced out for clarity. **B.** Structural overlay of Apo FhuA (PDB 1BY3, plug green) and of FhuA:Llp complex (PDB 8A60, plug pink, Llp purple), highlighting the conformational changes upon Llp binding. The N-terminus, plug loops 95–103 and 134–146, and EL7 and EL8 are displayed darker. Red arrows indicate the downwards displacement of plug loops upon Llp binding, and orange arrows the conformational changes of the extracellular loops EL7 and EL8. The breach in the upper part of the β-barrel between β-strands 12 (light blue) and 13 (plum) is labelled by a black double arrow. The front of the barrel has been removed for clarity. **C.** Structural overlay of soluble (gold) and FhuA-bound (purple) Llp. Conformational changes involving the flexible 39-44 loop (red arrow) and the C-terminus, including the FTE52–54 segment (circled red) are labelled.

Mutant DDK224–226-NNQ, located on periplasmic turn T2, shows partial protection and appears to have a lower binding affinity for Llp (Fig. 5). This is not unexpected since these residues are involved in the interactions with Llp, as seen in the FhuA-Ac-Llp structure and suggested by the related PISA analysis. The I9P and T12P mutants will be discussed below.

### 3. Analysis of the Llp mutants

We solved the structure of Llp in its free form and identified Llp residues perturbated upon FhuA addition by NMR (Fig. 3C). Based on these results, we designed a series of Llp mutants to probe the molecular determinants of the interaction with FhuA, prior to the release of the FhuA:Ac-Llp structure. Figure 4 recapitulates the T5 protection phenotype of Llp mutants. We analyse these mutants in the context of the now available FhuA:Ac-Llp structure. Llp mutant KD7-8AR behaves similarly to WT Llp with high FhuA protection, consistent with the structural data showing that these residues do not contribute to the FhuA-binding surface (Fig. 4C). The chemical shift perturbations observed by NMR upon FhuA addition likely result from altered interaction with detergent molecules bound to FhuA rather than direct contact with FhuA.

Llp mutants YK22-23AS, RK29-30SS and W46S show a mild loss of protection. Whereas it is not surprising for mutant YK22-23AS, as the mutated residues are not located at the interface with FhuA, it is surprising for mutants YK22-23AS and W46S, as K23 and W46 forms hydrogen bonds with FhuA P42 and N583 respectively. As the FhuA:Llp interaction surface is large, a point mutation could not be sufficient to strongly perturb it. A more substantial reduction of protection is observed for Llp mutant D28K. D28 is not directly involved in the interaction interface, but may create steric hindrance with FhuA residue F625 upon substitution. Llp triple mutant D18N-D21N-K23Q exhibits weak partial protection, indicating additive effects from multiple perturbations. In addition to the above-mentioned disruption of K23 bond with FhuA-P42, the D18K substitution likely abolishes two salt bridges with FhuA R297. The most affected Llp mutants are mutants YYMQD14–18AFAAK and YGSFW42–46STRAS, the mutated regions of both lying directly within the Llp:FhuA interface and involving more than one residue. Indeed the Llp 14–18 segment is involved in three hydrogen bonds with FhuA (between Llp Y17 and FhuA Q44 and K45, and between Llp D18 and FhuA R297). The Llp residues in the 42–46 segment form four hydrogen bonds with FhuA residues (Llp Y42, S44, and W46 with FhuA E56, E57, T54 and N583, respectively), highlighting the importance of the 39-44 loop in stabilising the complex (Fig. 6C). Interestingly, the FTE52-54YKK mutation reduces significantly the protection of *E. coli* against T5 infection, despite not being part of the direct contact interface. They probably affect the proper folding of the protein, as the mutated residues point towards the core of the protein, but HSQC analysis also reveals that these residues undergo significant chemical shift perturbations upon FhuA binding, suggesting an indirect structural role in FhuA binding. To compare the bound and unbound Llp structures, we calculated theoretical NOEs for the bound Llp structure and compared them to experimental NOEs of unbound Llp. Different NOE patterns were observed for residues D8-A13 and E54-F60, suggesting that these segments adopt different conformations in solution and in the FhuA-bound state. Superimposition of free Sol-Llp and bound Ac-Llp structures shows that it is indeed the case. It shows also that FhuA binding locks the 39-44 loop, flexible in free Llp, at the interface with FhuA (Fig. 6C). Analysis of Llp mutant FTE52-54YKK that shows only partial protection, while the mutated residues are not at FhuA interface, suggests that these residues are important for Llp conformational rearrangements, likely involving allostery.

### 4. Kinetics of Ac-Llp:FhuA formation

Two extreme mechanisms can be described in the formation of a protein-protein complex: conformational selection (A ⇌ A* ; A* + B ⇌ AB), where a rare pre-existing conformation is trapped by the interaction, or induced fit (A + B ⇌ AB* ⇌ AB), where rearrangements triggered by binding drive the formation of the final complex (Vega et al., 2016; Yu and Koshland, 2001).

The formation of the FhuA:Llp complex was assessed by different approaches. NMR allowed to identify key residues in the interaction, which roles were confirmed and refined by mutagenesis and in view of the FhuA:Llp structure. Other biophysical methods (MST, SPR, BLI, AUC and SEC) also allowed to evidence FhuA:Llp interaction in fast equilibrium and to measure a K_d_ in the µM range; however, a second slower interaction with a K_d_ an order of magnitude lower was also characterised, after longer incubation times. The complex formed after the first equilibrium does not lead to FhuA inactivation, and its formation is favoured by Llp acylation, which likely restrains diffusion of both proteins in the 2D membrane plane (or in the micellar phase). FhuA inactivation by Llp only occurs after the second equilibrium, strongly suggesting a significant contribution from induced fit. As described above, Llp binding requires the displacement of the periplasmic surface of FhuA plug (residues 20-51) (Fig. 6 A,B), which most probably happens during the second equilibrium. Interestingly, part of this plug surface (residues 20-31) is also remodelled following extracellular Fe-ferrichrome ligand binding, with the unwinding of the switch helix allowing binding of TonB, notably to the TonB box (Pawelek et al., 2006) (Fig. 6A). *In vivo*, Fe-ferrichrome binding probably initiate/facilitate these extensive structural rearrangements, which are associated with a high energetic barrier *in vitro*, and a strong temperature dependence of the kinetics (days at 4°C *vs*. hours at 37°C). As *i)* TonB binding partially occludes FhuA periplasmic surface, preventing Llp binding, *ii)* the FhuA:Llp complex forms *in vitro*, and *iii)* the FhuA T12P mutant, completely impaired in ligand transport, confers extra T5 protection by Llp binding, TonB-dependent transport does not seem to be required for Llp binding. In this context, the difference in Llp protection of FhuA mutants I9P and T12P, both located in the -disordered-TonB box is intriguing: while in our hands both mutants do not transport colicin M, the FhuA I9P mutant is not protected by Llp, while the FhuA T12P mutant is more protected by Llp than even the WT (Fig. 5B). *In vivo*, in the FhuA I9P mutant, the reorganisation of FhuA 20-51 segment, or Llp binding, could be prevented by an unproductive but strong interaction of FhuA TonB box with TonB. Inversely, in the FhuA T12P mutant, the reorganisation of the 20-51 segment could be facilitated by a much lower affinity of TonB to FhuA TonB box. *In vitro*, the absence of TonB would allow Llp binding to the I9P mutant.

## Conclusion

We present the structure of phage T5 protein Llp and analyse the complex it forms with FhuA. Acylation is essential for creating an active complex *in vitro* and probably *in vivo*. Analysis of Llp and FhuA mutants highlights the importance of the interface residues on the two proteins and of more distal regions that stabilise the complex, underscoring the importance of allostery, in both proteins, in Llp-mediated inhibition. Formation of the functional complex also involves a high activation energy, needed to displace residues 20-51, which form the periplasmic surface of FhuA plug. However, TonB-mediated iron transport is not necessary to form a functional complex.

## Material and methods

### 1. Molecular biology

We designed two constructs: the wild type Llp, with a His-tag in C-terminus, referred to as Ac-Llp, and a Maltose Binding Protein (MBP) fusion protein with Llp lacking its N-terminal cysteine to prevent acylation, termed MBP-Llp. The cleavage of the MBP moiety after purification results in a soluble form of Llp, Sol-Llp. Cloning of *Ac-llp* and *MalE-llp* genes were performed by PCR from T5 genomic DNA adapting the QuickChange Lightning Site-directed Mutagenesis Kit protocol (Agilent Technologies) to insert the genes into pET20b and pMAL-p2E plasmid, respectively. For *MalE-llp* a TEV site was inserted between the MBP sequence present on the pMAL-p2E plasmid and *llp*. *Ac-llp* was sub-cloned into pASK-IBA3C using QuickChange Lightning Site-directed Mutagenesis Kit to enable co-expression of the two proteins using the plasmid native *E. coli* promoter instead of a T7 promoter. All FhuA and Ac-Llp mutants were obtained using the same kit, and designed before the structure of FhuA:Llp was published.

### 2. Protein overexpression and purification

FhuA was produced as in (Flayhan et al., 2012), under the control of its natural promoter in the AW740 *E. coli* strain from a plasmid encoding the *fhuA* gene with a His-tag coding sequence inserted in the L5 extracellular loop. Cells were grown overnight in Lysogeny Broth (LB) medium at 37 °C in the presence of 125 µg.mL^-1^ ampicilin, 10 µg.mL^-1^ tetracyclin and 100 µM 2,2’ bipyridyl, an iron chelator inducing FhuA over-expression. After clarification of the cell lysate, total membranes were recovered by ultracentrifugation at 35,000 rmp for 45 min at 4°C (Ti45 rotor Beckman), and solubilised in 50 mM Tris pH 8.0, 2% (w/w) OPOE (N-octylpolyoxyethylene) at 37°C for 30 min. The insoluble material was recovered by ultracentrifugation in same condition as above and solubilised 1 h, 180 rmp at 37°C in 50 mM Tris pH 8.0, 1 mM EDTA, 1% (w/w) LDAO (N,N dimethyl dodecylamine-N-oxide, Anatrace). The solubilised fraction was supplemented with 4 mM MgCl_2_ and 5 mM imidazole and loaded onto a nickel affinity column (HiTrapTM Chelating HP 5 mL, GE Healthcare) previously equilibrated with 0.1 % LDAO, 20 mM Tris pH 8.0, 200 mM NaCl and washed with the same buffer. The protein was eluted from the column with 0.1% LDAO, 20 mM Tris pH 8.0, 200 mM imidazole, and loaded onto an ion exchange column (HiTrapTM Q HP 1 mL, GE Healthcare) equilibrated with 0.05% LDAO, 20 mM Tris pH 8.0. The protein was eluted by a 0 – 1 M NaCl linear gradient.

All interaction experiments were performed after detergent exchange from LDAO to DDM (Dodecyl-β-D-Maltopyranoside). FhuA was precipitated by a 10-time dilution into deionised water and recovered by centrifugation at 100,000 *g* for 45 min at 4°C. The pelleted protein was rinsed with deionized water and re-suspended at 1 mg.mL^-1^ into 25 mM Tris pH 8.5, and 1.3 % DDM.

Ac-Llp was produced in *E. coli* C43(DE3) strain in TB (Terrific Broth) supplemented with 125 µg.mL^-1^ ampicillin, at 20°C, without induction, for 24h. Cell lysis was performed in 50 mM Tris pH 8, 150 mM NaCl, 2 mM MgCl_2_, 1mg.mL^-1^ DNAse, 1X Clapa (1 µg.mL^-1^ Chymostatine, 1 µg.mL^-1^ Leupeptine, 1 µg.mL^-1^ Antipaine, 1 µg.mL^-1^ Pepstatine, 5 µg.mL^-1^ Aprotinine) by 10 passages in LM20 microfluidizer^TM^, at 14000 psi. Unbroken cells and large cellular debris were removed by low-speed centrifugation at 6,000 rpm for 15 minutes at 4 °C (JLA-8.1000 rotor, Beckman). Membranes were then collected by ultracentrifugation at 35,000 rpm for 20 minutes at 4 °C (Ti-45 rotor, Beckman). For solubilisation, membranes were adjusted to a final protein concentration of 2 mg.mL^-1^ in solubilisation buffer (25 mM Tris-HCl, pH 8.5, 150 mM NaCl, 4% OPOE) and incubated overnight at 4 °C with shaking. Insoluble material was removed by ultracentrifugation under the same conditions as described above. The supernatant was loaded onto a 5 mL HiTrap Chelating Nickel column (GE Healthcare), pre-equilibrated with 25 mM Tris pH 8.5, 150 mM NaCl, 0.3% OPOE. Detergent exchange was performed on-column, using a buffer containing 25 mM Tris pH 8.5, 150 mM NaCl, 1% DDM or LMNG or 2% OG (Octyl-β-Glucoside). Excess of detergent was washed away by the same buffer either containing 0.2% DDM or LMNG, or 0.64 % OG). Elution was performed using this buffer supplemented with 200 mM imidazole or 50 mM EDTA. Fractions containing Ac-Llp were pooled, diluted three times and loaded onto an anion exchange column (HiTrap Q HP 1 mL, GE Healthcare) pre-equilibrated with 25 mM Tris pH 8.5, 0.01% DDM. The protein was eluted using a 0–1 M NaCl linear gradient added to the equilibration buffer.

Sol-Llp was produced in *E. coli* BL21(DE3) strain in LB supplemented with 125 µg.mL^-1^ ampicillin at 30°C, with a 0.1 mM IPTG overnight induction. Lysis was performed as for Ac-Llp. A unique centrifugation at 100,000 g, for 20 min, at 4°C was performed and the supernatant was loaded onto a 5 mL MBP Trap™ HP Prepacked Column (GE Healthcare) pre-equilibrated with buffer A (25 mM Tris pH 8.5, 150 mM NaCl, 1 mM EDTA). Then the protein was eluted with 25 mM Tris pH 8, 150 mM NaCl, 1 mM EDTA, 10 mM maltose. The MBP was removed by TEV digestion using 1 mg of protease for 20 mg of MBP-Llp, overnight in buffer A supplemented with 3 mM reduced/0.3 mM oxidised Glutathione. The protein was then loaded onto a 5 mL HiTrap Chelating Nickel column (GE Healthcare). The last purification step was a size exclusion chromatography performed on a 25 mL Superdex 75 column pre-equilibrated with 25 mM Tris pH 6.5, 150 mM NaCl.

For NMR experiments, Sol-Llp was produced as described above with some modifications. Overexpression was carried out in M9 minimal medium (50 mM Na_2_HPO_4_, 20 mM K_2_HPO_4_, 10 mM NaCl, 2 mM MgSO_4_, 0.1 mM CaCl_2_, 0.1 mM MnCl_2_, 50 µM ZnSO_4_, 50 µM FeCl_3_) supplemented with 125 µg.mL^-1^ ampicillin and vitamins (1 mg.L^-1^ pyridoxine, biotin, hemi-calcium panthothenate, folic acid, choline chloride, niacin amide, 0.1 mg.L^-1^ riboflavin, 5 mg.L^-1^ thiamine), 4 g.L^-1^ ^13^C-glucose and/or 20 mM ^15^NH_4_Cl). Culture temperature was lowered down to 20°C before IPTG overnight induction. The final elution buffer pH was decreased down to 6.5.

To assess protein expression, 5 mL of bacterial culture were collected by centrifugation at 3,500 × *g*. Cell pellets were resuspended in 500 µL of 1× Laemmli buffer and sonicated for 30 seconds (without cooling). Samples were loaded onto an 18% SDS-PAGE gel at a volume of 12.5 µL per OD_600nm_ unit. Transfer to a 0.2 μm nitrocellulose membrane was performed using the Trans-Blot® Turbo™ system (Bio-Rad) with the “Bio-Rad Mixed” program (7 min/1.3 A/25 V) and the associated buffer (25 mM Tris, 192 mM glycine pH 8.3, 20% MeOH). The membrane was saturated (30 min in 0.1% TBS-Tween 20, 5% milk). Anti-histidine antibodies (Sigma-Aldrich) coupled to HRP were used at a 1:5000 dilution. Membrane was revealed after a 1-hour incubation at room temperature, the membrane was washed for 15 minutes with 0.1% TBS-Tween 20 and developed using SIGMAFAST™ 3,3ʹ-Diaminobenzidine tablets for colorimetric detection.

1. 3. Phenotypic assay

The various *E.coli* strains (C43(DE3), BL21(DE3), F and AW740), expressing Ac- or Sol-Llp, were pre-cultured at 37°C, 180 rpm, overnight in the presence of antibiotics when necessary. Bacterial cultures were then grown at 37°C up to OD _600 nm_ = 0.5. Then 1.5 mL of soft LB Agar (5 g.L^-1^) was inoculated with 0.3 mL of the culture and spread onto a LB agar Petri dishes. 5 µL of T5 serial dilutions following incubation or not with FhuA or FhuA/Llp complex were spotted onto the inoculated soft agar. Plates were incubated overnight at 37°C. The titre was estimated by counting lysis plaques in the spotted areas.

### 4. Thermal denaturation (nano-DSF)

Melting Temperature (Tm) was measured with a Prometheus NT.48 instrument (NanoTemper Technologies). The Tm of the two forms of Llp and of the FhuA:Llp complex were measured in the absence or presence of β-mercaptoethanol. 10 µl of 0.2 mg.mL^-1^ unlabelled protein was loaded into a glass capillary (Cat. No. PR-C002, nanoDSF Grade Standard Capillaries). Unfolding curves were obtained for temperatures ranging from 20 to 95°C, with 1°C.min^-1^ ramp, at 20% excitation power. The Tm were calculated with PR.ThermControl software (NanoTemper Technologies) by analysing the first derivative curve of autofluorescence at 350 nm plotted against the temperature for Llp, or of the first derivative curve of the 350/330 nm ratio for the FhuA samples. Samples were incubated two days at room temperature before measurement.

### 5. Microscale thermophoresis (MST)

FhuA was labelled with a fluorophore using the Monolith His-Tag Labelling Kit RED-tris-NTA 2nd Generation (NanoTemper Technologies) according to the manufacturer protocol. FhuA concentration was set at 250 nM and the untagged proteins Ac-Llp, Sol-Llp or detergent micelles were serially diluted in the same buffer as FhuA (25 mM Tris pH8.5, 150 mM NaCl, 0.2 mM DDM). Sol-Llp was tested between 48 nM and 1.6 mM, Ac-Llp between 5.3 nM and 887 µM and DDM micelles between 0.383 and 196 mM. FhuA was titrated in a 1:1 dilution with Llp and inserted into Monolith NT.115 capillaries. The measurements were performed using a power of 30% for the LED (light-emitting diode) and 20% for the MST for 20 s. The K_d_ was determined using the MST analysis of the Initial fluorescence was normalised by the MO.Affinity software prior to fitting. The MST signal was found to be affected by detergent concentrations, therefore, to minimise detergent variation during titration, Ac-Llp dilutions were supplemented with DDM to maintain a constant total DDM concentration across the titration series.

### 6. Other interaction measurements

SPR (Surface Plasmon Resonance) was performed using CM3 series S sensor chip where FhuA was immobilised *via* its lysins by EDC/NHS treatment at a density of 1910 Response Unit (RU) on flow cell 2 (Fc2) and 3400 RU on flow cell 4 (Fc4). Ac-Llp or Sol-Llp was injected onto the chip at concentrations ranging from 0.68 to 700 µM and 0.99 to 509 µM, respectively. Surface was regenerated between injections with a 50 mM Glycine pH12, 25 mM EDTA, 2.3 mM TritonX100, 0.2 mM DMM or 0.1 mM LMNG (Lauryl maltose neopentyl glycol) regeneration buffer.

For BLI (Bio-layer Interferometry) experiments, FhuA and Sol-Llp proteins were biotinylated using EZ-LinkTM Sulfo-NHS-LC-Biotin kit (Thermo ScientificTM). FhuA or Sol-Llp biotinylated proteins were immobilized on Octet® SA Biosensors (Sartorius) for 600 s at 1000 rpm agitation. The protein concentrations tested during immobilization were 5 µg.mL^-1^, 10 µg.mL^-1^ and 50 µg.mL^-1^for FhuA and 1 µg.mL^-1^, 5 µg.mL^-1^ and 50 µg.mL^-1^ for Sol-Llp. For measurements, all steps were performed at 1000 rpm with an association and dissociation time of 600 s at protein concentration ranges similar to the ones used in SPR. Sensors were regenerated for 20 s at 1000 rpm in the same regeneration buffer than that for SPR. The sensorgrams were processed with the dedicated manufacturer’s software “Data Analysis HT 11.1 software”.

### 7. Near-UV CD

The measurements were carried out on a JASCO J-810 circular dichroism (CD) spectropolarimeter equipped with a temperature controller (Peltier system). The spectra were recorded on Ac-Llp and Sol-Llp proteins at 30 μM in 25 mM Tris pH 8.5, 150 mM NaF and 0.4 mM DDM for Ac-Llp, in the absence or presence of 100 mM EDTA. The spectra were recorded over 15 successive acquisitions in a 1 cm cuvette. The values are valid for a voltage lower than 700 V, *i.e*. in our case between 260 nm and 200 nm (far UV). Values were normalised by blank subtraction and converted to mean molar ellipticity (deg.cm^2^.dmol^-1^) using the following formula:

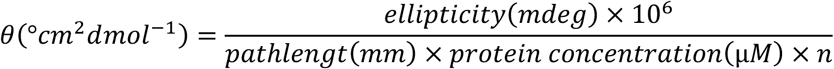

Where n is the number of peptide bonds in the protein. Secondary structure type prediction was performed using an online tool K2D (https://www.bork.embl.de/~andrade/k2d/).

### 8. AUC & SEC-MALLS

Sedimentation velocity AUC (SV-AUC) experiments were conducted in an XLI analytical ultracentrifuge (Beckman, Palo Alto, CA) using an An-60Ti or An-50Ti rotor, using double channel centre pieces (Nanolytics, Germany) of 12, 3 or 1.5 mm optical path length equipped with sapphire windows, with the reference channel being typically filled with solvent without detergent. Acquisition was done at 20°C and at 42,000 rpm (130,000 *g*), overnight, using absorbance (280 nm) and interference detection. Data processing and analysis was done using the program SEDFIT (Schuck, 2000) from P. Schuck (NIH, USA), REDATE (Zhao et al., 2015) and GUSSI (Brautigam, 2015). Corrected s-values for solvent density and viscosity, s_20w_, are given in the text. They were calculated, for membrane proteins, considering for their partial specific volume (*v̄*), the mean value considering 1g of bound detergent per g of protein.used were *v̄* 0.725 mL.g^-1^ (FhuA), 0.734 mL.g^-1^ (Ac-Llp) and 0.725 mL.g^-1^ (Sol Llp).

SEC-LS experiments were conducted on a HPLC equipped with UV detector (Shimadzu) and light scattering and differential refractive index detectors (Wyatt). FhuA and Ac-Llp or sol-Llp samples were analysed independently and in complex. Separation was performed on a Superdex 200 10/300 GL column (GE Healthcare) equilibrated at 20°C with an elution buffer containing 20 mM Tris (pH 8.5), 150 mM NaCl and 0.2% DDM. Data were analysed using protein conjugate method with the ASTRA 5 software. Analyses parameter used : dn/dc, epsilon was 0.191 mL.g^-1^, 1.385 mL.mg^-1^.cm^-1^ (FhuA), 0.197 mL.g^-1^, 1.659 (Ac-Llp) and 0.197 mL.g^-1^, 1.857 mL.mg^-1^.cm^-1^ (Sol Llp).

### 9. NMR

NMR measurements were carried out on 0.950 mM uniformly ^15^N- or ^15^N- and ^13^C-labelled sample of Sol-Llp in 90% H_2_O (25 mM Tris pH 6.5, 75 mM NaCl) and 10% D_2_O. The NMR spectra were acquired at 14.1 T and 23.5 T (600 and 1000 MHz of proton Larmor frequency, respectively) on Bruker Avance III spectrometers, all equipped with a cryogenic cooled 5 mm TCI probehead. NMR measurements were carried at 303 K. Sample state was checked using [^1^H-^15^N] HSQC spectra and methyl-TROSY spectra of Ac-Llp and Sol-Llp in natural abundance of isotopes.

The assignment of Llp backbone spin system was performed at 14.1 T, using the following set of 3D triple-resonance experiments: HNCO, HNcaCO, HNCA, HNcoCA, HNcaCB, HNcocaCB as well as standard CBCACONH (Hiller et al., 2008), HACACONH (Fiorito et al., 2006), and HACANH (Hiller et al., 2005) APSY experiments. The HA and HB protons were assigned with HAHBNH and HAHBcoNH spectra measured at 23.5T. Additional ^15^N-resolved [^1^H,^1^H]-NOESY spectrum and two 3D ^13^C-resolved [^1^H,^1^H]-NOESY spectra with carrier frequencies in the aliphatic and aromatic regions, respectively, were measured at 23.5T for side spin system assignment and structural restraint acquisition.

#### Spin system assignment and structure calculation

Sequence-specific backbone resonance assignments were independently determined by manual and automated analysis of the 2D [^1^H-^15^N] HSQC and the six 3D triple resonance experiments mentioned above. Manual peak picking, spin system generation and sequence-specific assignment were performed using the program CCPNmr 3 (Skinner et al., 2016). The input for automated backbone resonance assignment using the program UNIO-MATCH version 2.9.5 (Volk et al., 2008) consisted of the protein sequence and manually generated lists of peak positions in the six triple-resonance spectra. The manual and automated approaches produced identical results for the backbone assignments.

The input for automated side chain assignment using UNIO-ATNOS/ASCAN (Fiorito et al., 2008) consisted of the protein sequence, the list of NMR chemical shifts from the previous backbone assignment, and the three 3D NOESY spectra.

The input for automated NOESY signal identification (peak picking) and NOE assignment using UNIO-ATNOS/CANDID (Guerry et al., 2015) consisted of the UNIO-MATCH chemical shift assignments for the polypeptide backbone, the UNIO-ATNOS/ASCAN output of side chain chemical shift assignments, and the three NOESY spectra. The standard UNIO protocol was employed. The final structural bundle was energy-refined in explicit solvent using the program CNS_SOLVE version 1.3 (Brunger, 2007) employing the standard procedure recommended by the RECOORD database initiative (http://www.ebi.ac.uk/pdbe/recalculated-nmr-data) (Nederveen et al., 2005).

#### Structure validation

The analysis of the stereochemical quality of the molecular models was performed using the PDB validation tools (https://validate-rcsb-3.wwpdb.org) (Gore et al., 2017).

#### NMR for probing interaction

[^1^H,^15^N]-SOFAST-HMQC was recorded to investigate the detergent influence on the lipoprotein and the complex formation with FhuA. Chemical shifts and peak intensity variations were followed with serial addition of Llp to increase [Llp]/[FhuA] ratio. The samples were mixed using stocks of Llp at 1.2 mM and FhuA at 0.15 mM, see table S5B.

#### Dynamics measurements

^15^N spin relaxation measurements were performed using well established R_1_, R_1ρ_, and {^1^H}-^15^N nOe experiments (Lakomek et al., 2012). Magnetisation decays were set in randomized order to 0, 80, 160, 240, 320, 400 (x2), 560, 720, 880 and 1120 ms for R_1_ and 1, 10, 20, 40, 60 (x2), 90, 120, 160, 200, 250, ms for R_1ρ_. R2 rates were obtained using R_2_ = R_1ρ_ / sin^2^ θ - R_1_ / tan^2^θ where tan θ = ω_1_/Ω, ω_1_ being the spin-lock field strength and Ω the offset from the 15N carrier. For the measurement of the nOe, a recycling delays of 5 s was used and 3s of amide proton saturation was performed using a series of (Δ-180-Δ). The FUDA package was used to extract relaxation rates. Model-free analysis of Llp was performed using TENSOR2 (Dosset et al., 2000).

### 10. In silico protein analysis

To predict the structure of proteins, the ColabFold notebook, whose structure prediction is powered by AlphaFold2 combined with a fast, multiple sequence alignment generation stage using MMseqs2 was used (Mirdita et al., 2022).

## Acknowledgment

We acknowledge the M1 internship work of Perrine Guenneau in the characterisation of the FhuA mutants, Christine Ebel for very useful discussions, Fabienne Devime and Stéphane Ravanel (LPCV, Grenoble) for ICP-MS measurements on Sol-Llp and Thierry Touzé (I2BC) for a kind gift of Colicin M.

This research was funded by the Agence Nationale de la Recherche, grant numbers ANR-21-CE11-0023 and ANR-22-CE92-0046-01 to CB. It used the biophysical platform at the Grenoble Instruct-ERIC Center (ISBG; UAR 3518 CNRS-CEA-UGA-EMBL) within the Grenoble Partnership for Structural Biology (PSB). IBS platform access is supported by FRISBI (ANR-10-INBS-05-02) and GRAL, a project of the University Grenoble Alpes graduate school (Ecoles Universitaires de Recherche,) CBH-EUR-GS (ANR-17-EURE-0003). Financial support from the IR INFRANALYTICS FR2054 is gratefully acknowledged. IBS acknowledges integration into the Interdisciplinary Research Institute of Grenoble (IRIG, CEA).

## Author Contributions

CB conceived the project. SD performed all the *in vitro* and *in vivo* experiments, with help from CDV and CD. CDV helped with the design and obtention of all mutants and the BLI and SPR experiments. AlR performed the AUC and the SEC-MALS experiments. CM helped with MST and Circular Dichroism. EM and SD performed NMR and analysed interaction and relaxation data, under the supervision of LS. SD and TH calculated the structure. SD and CB analysed and interpreted the data. SD wrote the first draft of the paper with the help of CDV. CB corrected the manuscript and all authors approved it.

## Data Availability

The ^1^H, ^13^C and ^15^N chemical shifts have been deposited in the BioMagResBank under the BMRB Id 34696. The atomic coordinates of bundles of 20 NMR Sol-Llp conformers are deposited in the Protein Data Bank under the accession code 7QJF.

