## Supplementary Figures for "Superinfection exclusion strategy of siphophage T5: analysis of the FhuA:Llp complex"

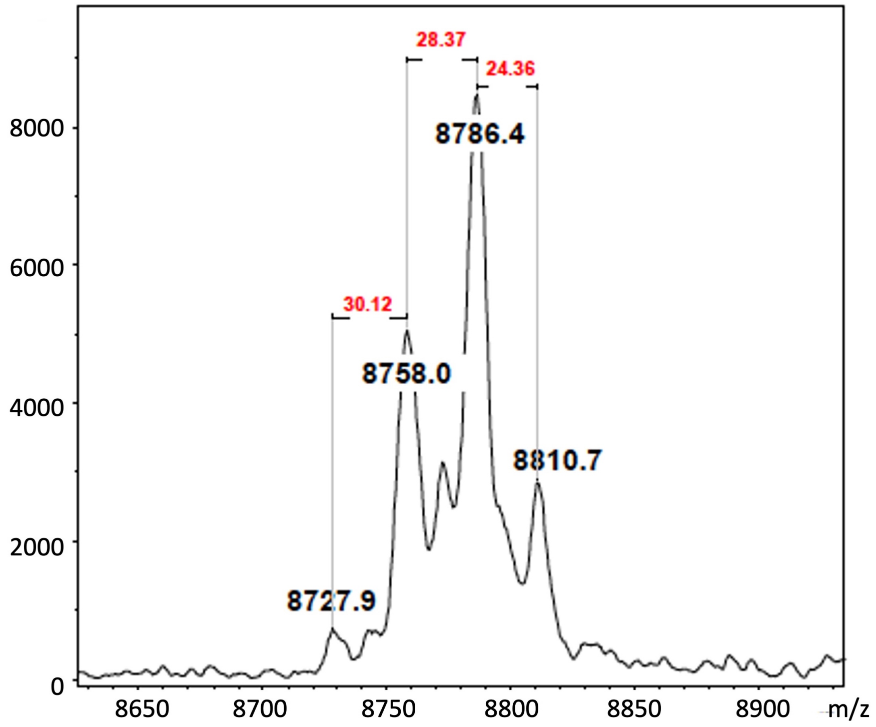


**Figure S1: Mass spectrometry analysis of purified Ac-Llp.** The x-axis represents the mass-to-charge ratio (m/z) and the y-axis the intensity (A.U.). The differences in m/z, in red, are compatible with the deletion/addition of a CH=CH, a $CH_2$-$CH_2$ and a $CH_2$-$CH_2$ and an unsaturation, for the 24, 28 and 30 Da values, respectively. The observed species are consistent with lipidation by fatty acid chains of varying length ($C_{12}$–$C_{18}$), reflecting the natural heterogeneity of *E. coli* membrane lipids.

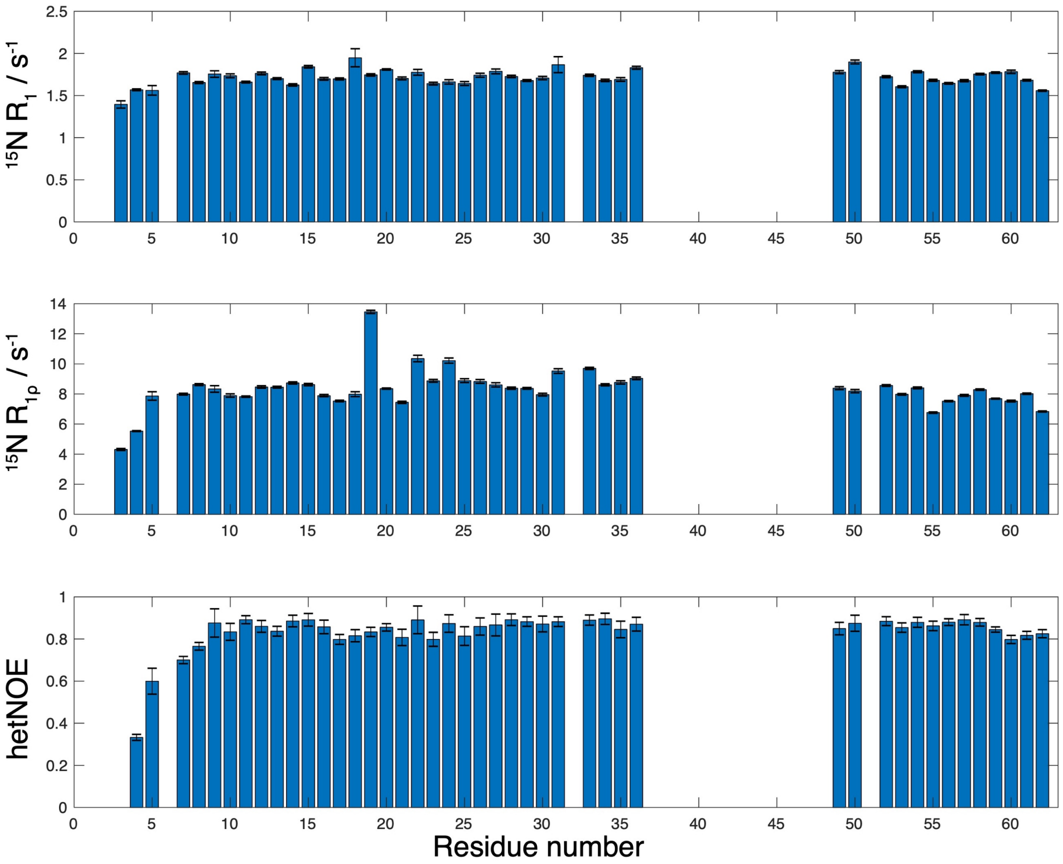


**Figure S2: NMR studies of Sol-Llp dynamics and interaction.** Residue-specific $^{15}N$ $R_1$ (top), $R_2$ (middle) and {$^1H$}–$^{15}N$ heteronuclear NOE (bottom) values measured for Sol-Llp. Error bars indicate standard deviations from replicate measurements. Missing data correspond to unassigned residues.

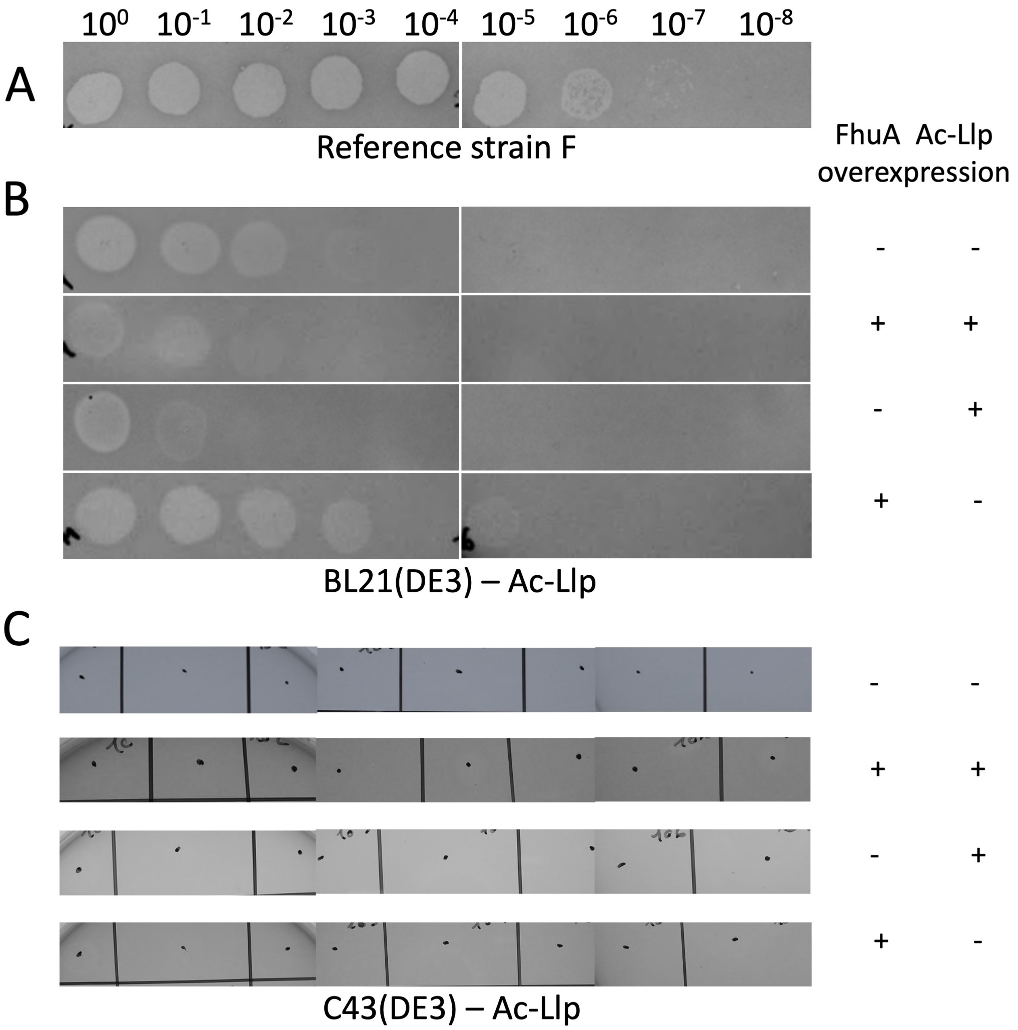


**Figure S3:** ***In vivo E. coli*** **T5 protection by Llp.** Serial T5 dilutions were plated on reference strain F, which displays the same infectivity as BL21(DE3), as a control (**A**), BL21(DE3) (**B)** and C43(DE3) (C), with basal (-) or overexpression (+) of FhuA and Ac-Llp by the iron chelator Dipyridyl at 0.3 mM and IPTG at 1 mM, respectively.

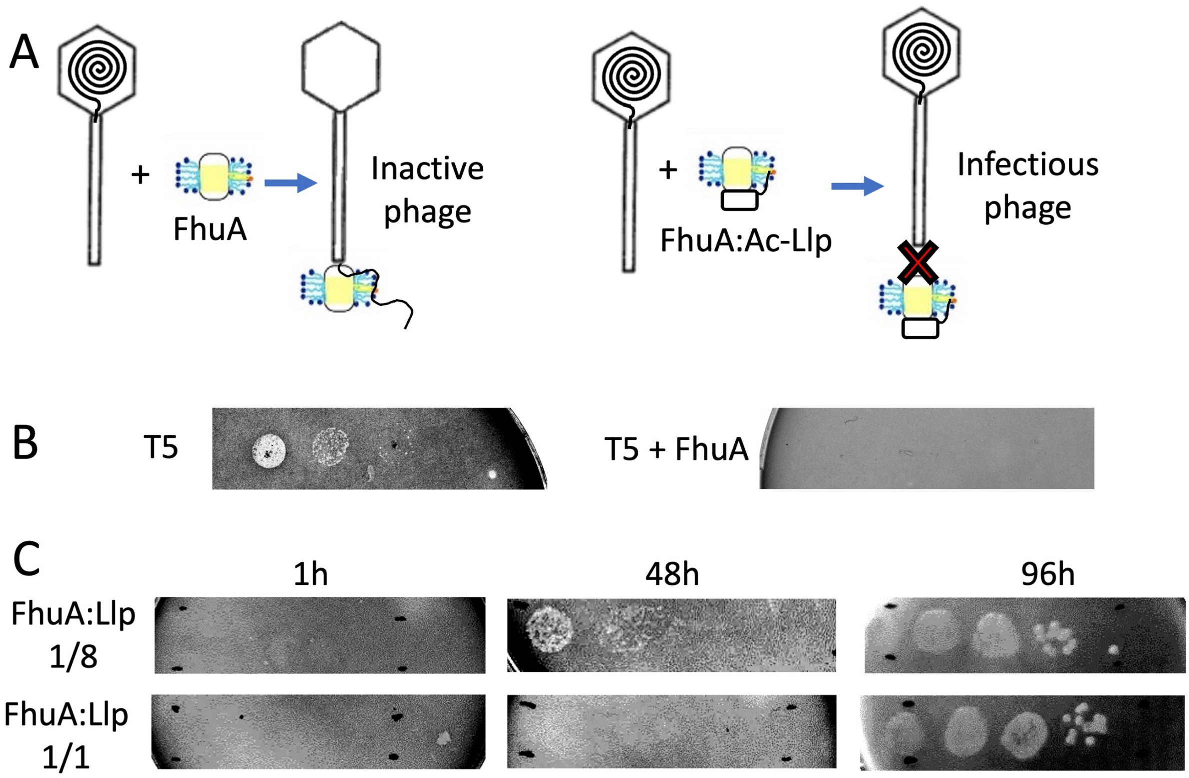


**Figure S4. *In vitro* investigation of the FhuA-Ac-Llp complex. A**. schematic illustration of the assay: T5 incubation with FhuA results in a non-infectious phage (left), whereas incubation with an active FhuA:Ac-Llp complex leads to an infectious phage (right). **B**. left: Serial dilutions of T5, plated on strain F, indicates an infectious phage up to a dilution at $10^{-3}$. Right: Serial dilution of T5 after incubation with FhuA (110 nM). T5 infectivity is completely abolished. **C**. Serial dilutions of T5 after incubation with FhuA:Ac-Llp at 1:8 or 1:1, at 4°C for 1, 48, or 96 h.

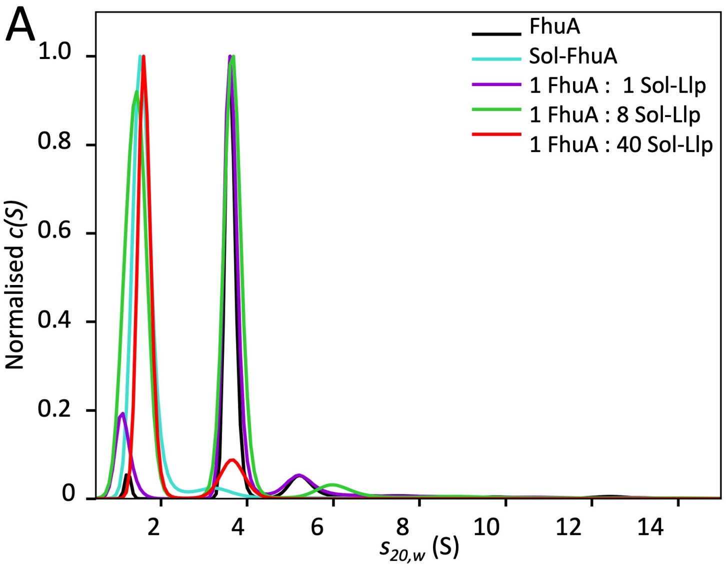


B

| [Llp] μM | [FhuA] μM | [Llp:FhuA] (%) |
|---|---|---|
| 46 | 127 | 0.4 |
| 105 | 120 | 0.9 |
| 166 | 114 | 1.5 |
| 232 | 106 | 2.2 |
| 231 | 75 | 3.1 |
| 280 | 71 | 3.9 |
| 331 | 67 | 4.9 |
| 387 | 63 | 6.2 |
| 436 | 59 | 7.4 |

**Figure S5: Characterisation of FhuA interaction with Sol-Llp by AUC and NMR. A**. C(s) distribution (SV-AUC) in absorbance of FhuA:Sol-Llp complexes at different molar ratios (colours are indicated in the figure), after 7 days of prior incubation at 4°C. FhuA was solubilised in 0.05% LDAO. The values of the sedimentation coefficients $s_{20,w}$, of 1.5 and 3.6 S, for Sol-Llp and solubilized FhuA, respectively, are the same in the different samples. **B.** NMR titration experiments were performed using $^{15}$N-labelled Sol-Llp and unlabelled FhuA for the different noted concentrations of Sol-Llp and FhuA. The percentage of complex formed was calculated assuming a dissociation constant ($K_d$) of 1.5 mM, as determined by MST.

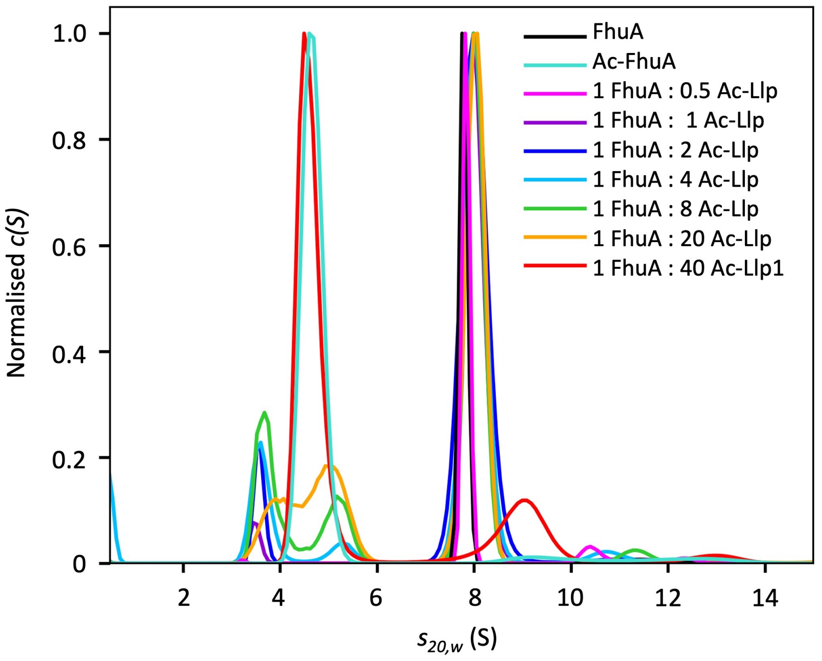


**Figure S6: Characterisation of FhuA interaction with Ac-Llp by AUC.** C(s) distribution (SV-AUC) in absorbance of FhuA:Ac-Llp complexes at different molar ratios (colours are indicated in the figure), after 7 days of prior incubation at 4°C. FhuA was solubilised in 0.01% DDM. DDM sediments with $s_{20\ w}$ at ≈3.5 S, Ac-Llp at 4.7 S, FhuA at 7.8 S. In the presence of Llp at up to Ac-Llp/FhuA ratios of 20, the sedimentation of FhuA increases to 8 S.

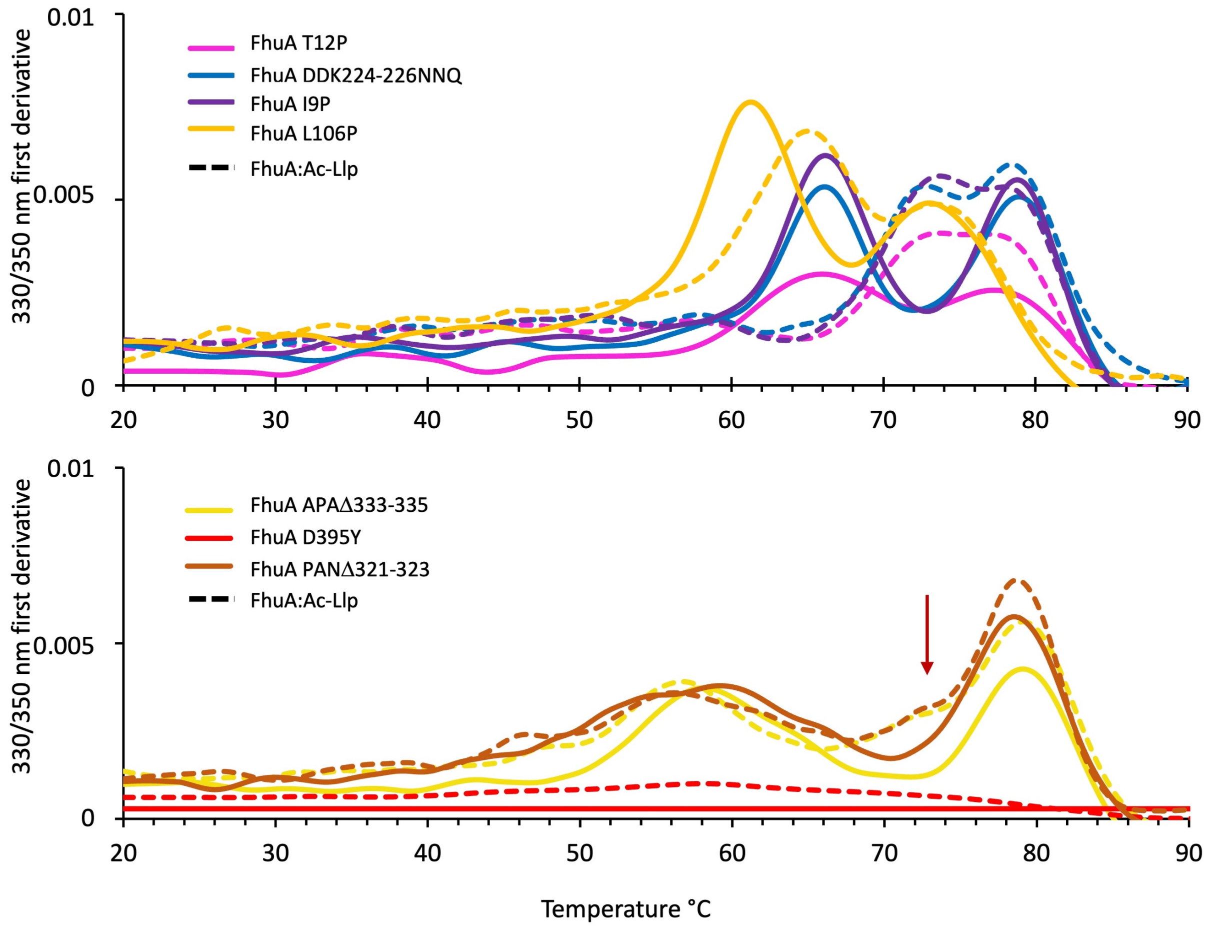


**Figure S7. Characterisation of FhuA mutant interactions with Ac-Llp by NanoDSF and MST. A.** Thermal denaturation profiles of FhuA mutants (solid line) and their corresponding FhuA:Ac-Llp complexes (dashed line) measured by NanoDSF. Different colours correspond to the different FhuA mutants as indicated in the figure. FhuA mutants with two clear transitions (plug and β-barrel Tm) are shown in the upper panel, while those without a clear plug transition are shown in the lower panel. The red arrow points to the shoulder that aligns with the plug denaturation maximum in the WT FhuA:Ac-Llp curve.

A

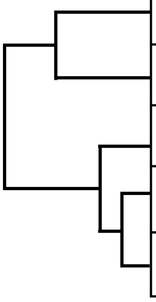

| | T5 -Llp | T1-Llp | φ80-Cor | HK022-Cor |
|---|---|---|---|---|
| T5 -Llp | | | | |
| T1-Llp | 23 | | | |
| φ80-Cor | 21 | 37 | | |
| HK022-Cor | 19 | 36 | 79 | |
| mEp167-Cor | 20 | 35 | 77 | 90 |

<25 | 25-50 | 50-80 | >80

B

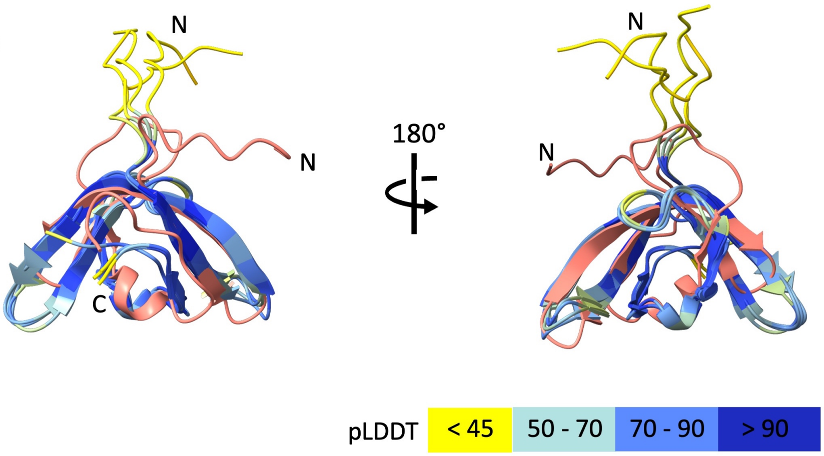


**Figure S8. Structural homology within the Cor and Llp families despite low sequence identity. A.** Colour-coded pairwise sequence identity matrix derived from Clustal Omega alignment of full-length T5 and T1 Llp and Cor proteins (81 aa). **B.** Structural superposition of the NMR structure of Sol-Llp (salmon) with AlphaFold-predicted models of the above-mentioned Cor and Llp proteins, coloured according to per-residue confidence (pLDDT score).

| Quantity | Value |
| --- | --- |
| NOE-derived upper distance restraints[a] | 511 |
| Intraresidual (\|i – j\| = 0) | 130 |
| Short-range (\|i – j\| = 1) | 140 |
| Medium-range (2 ≤ \|i – j\| ≤ 4) | 62 |
| Long-range(\|i – j\| ≥ 5) | 179 |
| Dihedral angle constraints | 106 |
| Residual target function[b], Å$^2$ | 1.53 ± 0.37 |
| Residual NOE violations | |
| Number > 0.2 Å | 0 |
| Residual angle violations | |
| Number > 5 ° | 0 |
| Amber energies, kcal/mol | |
| Total | -1909.7 ± 69.3 |
| Van der Waals | -215.5 ± 12.5 |
| Electrostatic | -2268.3 ± 69.5 |
| RMSD from ideal geometry | |
| Bond lengths, Å | 0.0158 ± 0.0007 |
| Bond angles, ° | 1.80 ± 0.09 |
| RMSD to the mean coordinates[c], Å | |
| bb | 1.02 ± 0.19 |
| ha | 1.58 ± 0.2 |
| Ramachandran plot statistics[d] | |
| Most favored regions (%) | 66.7 |
| Additionally allowed regions (%) | 28.5 |
| Generously allowed regions (%) | 3.1 |
| Disallowed regions (%) | 1.7 |

**Table S1: Inputs for structure calculation and NMR structure characterization of the energy-minimized Sol-Llp structure.** [a]Number of meaningful, non-redundant upper distance restraints between residues with sequence positions *i* and *j*. [b]Average residual target function value of all 20 structures calculated by CYANA. [c] 'bb' indicates backbone atoms N, C, Cα and C'; 'ha' indicates all heavy atoms. Root-mean-square deviations (RMSD) are reported for the well-defined structural regions of the protein, which include residues 6–38 and 45–62. [d]Ramachandran plot statistics were determined using the program PROCHECK (Laskowski et al., 1993).
